# Senescence: Still an Unsolved Problem of Biology

**DOI:** 10.1101/739730

**Authors:** Mark Roper, Pol Capdevila, Roberto Salguero-Gómez

**Affiliations:** Department of Zoology, University of Oxford, 11a Mansfield Road, Oxford, OX1 3SZ, UK; Centre for Biodiversity and Conservation Science, University of Queensland, St Lucia 4071 QLD, Australia; Evolutionary Demography laboratory, Max Plank Institute for Demographic Research, Rostock 18057, Germany

## Abstract

Peter Medawar’s ‘An Unsolved Problem of Biology’^1^ was one of several formal attempts to provide an explanation for the evolution of senescence, the increasing risk of mortality and decline in reproduction with age after achieving maturity. Despite *ca.* seven decades of theoretical elaboration aiming to explain the problem since Medawar first outlined it, we argue that this fundamental problem of biology remains unsolved. Here, we utilise demographic information^2, 3^ for 308 multicellular species to derive age-based trajectories of mortality and reproduction that provide evidence against the predictions of the classical, still prevailing, theories of ageing^1, 4, 5, 6^. These theories predict the inescapability of senescence^1, 4^, or its universality at least among species with a clear germ-soma barrier^5, 6^. The patterns of senescence in animals and plants that we report contradict both of these predictions. With the largest ageing comparative dataset of these characteristics to date, we build on recent evidence^7, 8^ to show that senescence is not the rule, and highlight the discrepancy between existing evidence and theory^7, 8, 9^. We also show that species’ age patterns of mortality and reproduction often follow divergent patterns, suggesting that organisms may display senescence for one component but not the other. We propose that ageing research will benefit from widening its classical theories beyond merely individual chronological age; key life history traits such as size, the ecology of the organism, and kin selection, may together play a hidden, yet integral role in shaping senescence outcomes.

## Main

The evolution of senescence has long been explained by a collation of theories defining the ‘classical evolutionary framework of ageing’. The central logic common to the theories argues that the force of natural selection weakens with age^1, 4, 5, 6^. Selection becomes too weak to oppose the accumulation of genes that negatively affect older age classes^1^, or favours these genes if they also have beneficial effects at earlier ages in life^5^, when the contribution individuals make to future populations is assumed to be greater. Selection, therefore, should favour resource investment into earlier reproduction rather than late-life maintenance^6^. Ultimately, these theories predict, directly^4^ or indirectly^1, 5^, that senescence is inescapable^4^, or at least inevitable in organisms with clear germline-soma separation^5, 6^.

It takes one example to disprove any rule. Recently, a comparative description of demographic patterns of ageing in 46 species of animals, plants, and algae^8^ contradicted the predictions of the classical evolutionary framework. Many of the examined species displayed negligible^10^ or even negative^11^ senescence, where the risk of mortality remains constant or decreases with age, and reproduction remains constant or increases with age. This mismatch between observations and expectations have rendered the classical evolutionary framework insufficient to explain the evolution of senescence across the tree of life. We now need to understand mechanisms behind such variation of ageing patterns^9^, and how prevalent such “exceptions” are to the rule of ageing.

Here, we utilise high-resolution demographic information for 48 animal^2^ and 260 plant^3^ species worldwide to (i) provide a quantitative evaluation of the rates of actuarial senescence – the progressive age-dependent increase in mortality risk with age after maturation – across multicellular organisms, (ii) test whether the classical evolutionary framework explains the examined diversity of senescence rates, with special attention to predictions from germ-soma separation, and (iii) propose how to widen the classical evolutionary framework of ageing to better encompass the tree of life.

First, we derived life tables^12^ from a selection of species’ matrix population models^13^, each of which are a summary of the population dynamics of the species in question under natural conditions (See Methods & Supplementary Information). We then quantified the rate of actuarial senescence on the survivorship trajectory of each species’ life table using Keyfitz’ entropy (*H*)^14^. This metric quantifies the spread and timing of mortality events in a cohort as individuals age, with *H* <1 indicating that most mortality occurs later in life (*i.e.* actuarial senescence), and *H*>1 indicating low mortality at advanced ages, whereby individuals may escape actuarial senescence (See Methods & Supplementary Information). Importantly, *H* is normalised by mean life expectancy, facilitating cross-species comparison to examine general patterns and plausible mechanisms.

**Figure 1.**
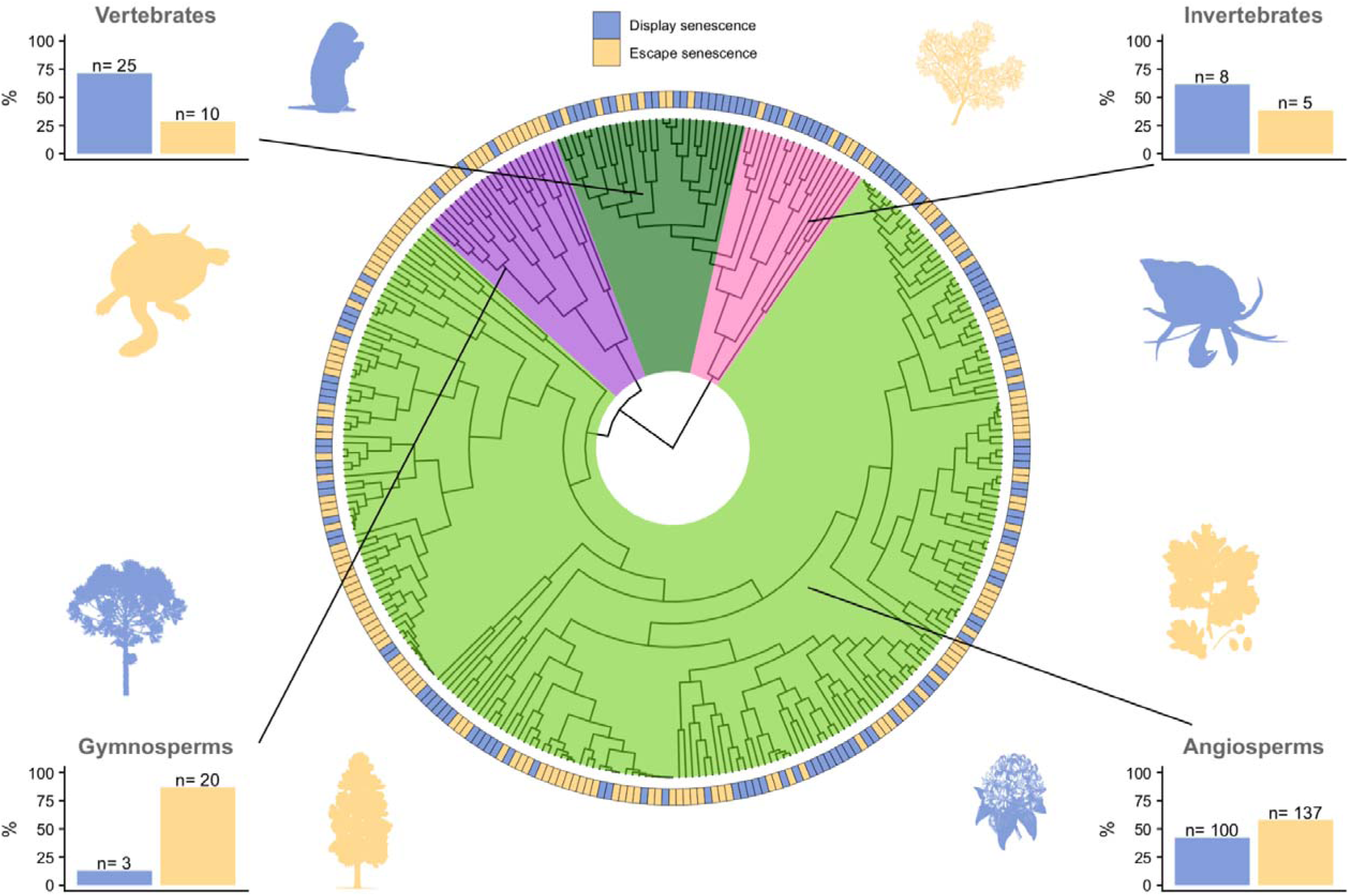
The evolution of and escape from senescence across multicellular life. The classical evolutionary framework of ageing does not explain the evolution of actuarial senescence across our 308 study species. Species that escape (yellow) and display (blue) senescence, as measured by Keyfitz’ entropy (H)14, are dispersed throughout the four examined clades, with the percentages of each clade either displaying or escaping actuarial senescence shown in the bar charts. Depicted around the phylogeny are eight representative species, escaping (blue) and displaying (yellow), from each clade. Clockwise, these species are Paramuricea clavata (Invertebrate), Pagurus longicarpus (Invertebrate), Quercus rugosa (Angiosperm), Rhododendron maximum (Angiosperm), Pinus lambertiana (Gymnosperm), Taxus floridana (Gymnosperm), Marmota flaviventris (Vertebrate) and Chelodina expansa (Vertebrate).

Most of the species in our analysis do not display senescence (Fig. 1, Supplementary Table 1). These include approximately 30% (10/35) of the studied vertebrate species, which have a clear germ-soma separation, such as the broad-shelled river turtle (*Chelodina expansa;* Fig. 1). In contrast, 42% of vascular plants (angiosperms and gymnosperms), all lacking a clear germ-soma barrier, display senescence (Fig. 1). In general, the evolutionary history of a species in our study plays a relatively weak role in constraining its ability to escape/evolve senescence. Estimates of Pagel’s λ^15^, a metric that measures how well phylogenetic relatedness predicts the variation of a trait across species (See Methods and Supplementary information) are generally weak (Extended Data Table 1), with the full analysis producing a Pagel’s λ = 0.38 (0.14-0.65, 95% C.I.). Overall, the emerging senescence landscape appears (i) (i) not inescapable, (ii) not inevitable in species with a germ-soma barrier, and (iii) prevalent in species without a clear germ-soma barrier. These findings are in direct contradiction with the predictions of the classical evolutionary framework of ageing^1, 4, 5, 6^.

The central assumption of the classical evolutionary framework of ageing, that the force of natural selection weakens with age, rests on the assumption that older individuals contribute less to future populations^1, 4, 5, 6^. This is both because fewer individuals survive to later age classes^1^, and individuals are expected to favour reproduction at young rather than old ages^1, 5, 6^. To observe how different age classes contribute to future populations in our study species, we use the derived life tables^12^ to quantify age-specific reproduction rates (*m(x)*) for species that we previously identified to display actuarial senescence (*H*<1) *vs.* those that do not (*H*>1) (See Methods & Supplementary information). The *m(x)* trajectories for all the species in our dataset can be found in Extended Data Fig. 1. For practicality, here, we provide the trajectories for the eight representative species across the four clades depicted in Figure 1.

Classical theories of ageing predict that rates of reproduction should decline with age,^1, 4, 5, 6^. In our study species, however, patterns of *m(x)* are diverse and appear to be independent of whether the species displays or escapes actuarial senescence (Extended Data Fig. 1a-h). For instance, the long-wristed hermit crab (*Pagarus longicarpus*) displays actuarial senescence (Fig. 1), yet its reproduction increases with age (Fig. 2). Our results suggest that an individual’s risk of mortality and rate of reproduction often follow different trajectories (Fig. 2, Extended Data Fig. 1a-h). For each species in our study, we only consider a single studied population, and so this decoupling cannot be an artefact of intra-specific variation across different populations. It follows that species may display actuarial senescence, but not reproductive senescence, and *vice versa*. Thus, we urge future work to consider that senescence is a two-component phenomenon of which, as displayed here, both are not destined to the same fate. To fully divulge the senescence profile of a species one must consider both mortality and reproduction.

**Figure 2.**
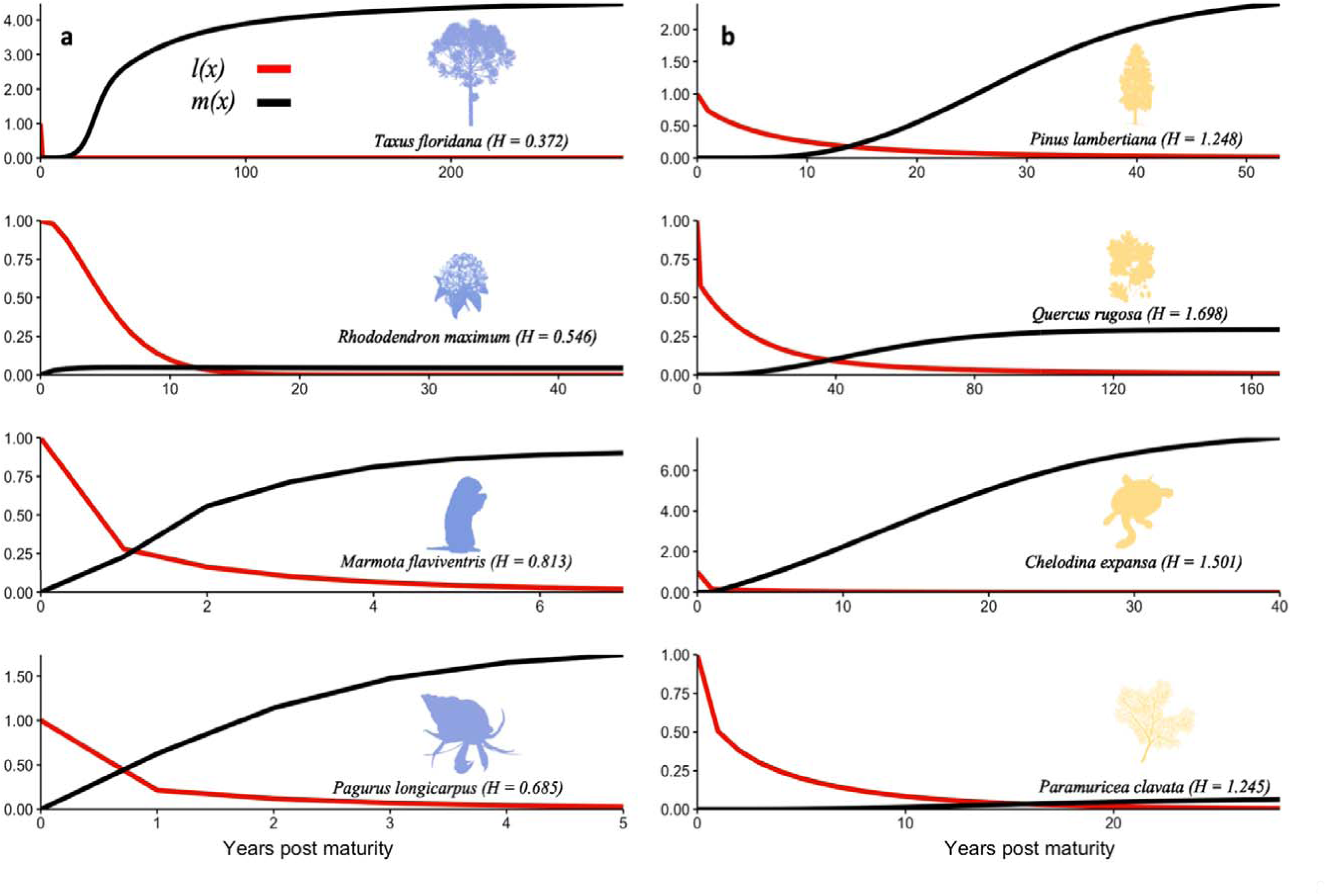
Age-based patterns of survivorship (l(x) - red) and reproduction (m(x) - black) are often decoupled, as shown for a selected subset of the examined species in Figure 1. **(a)** Species quantified as displaying actuarial senescence (H < 1) and **b)** species that escape actuarial senescence (H > 1). Species are representative of vertebrates, invertebrates, gymnosperms, and angiosperms. Trajectories are conditional upon reaching the age of maturity, represented as 0, at which the mature cohort is defined to have entered adulthood with a survivorship of 1. The trajectories of l(x) and m(x) run from age at maturity to the age at which 5% of the mature cohort is still alive.

Studies on reproductive senescence are sparser than their actuarial senescence counterparts. Although, important longitudinal investigations into reproductive senescence have been conducted^16, 17, 18^, and current data suggest that rates of reproduction, like mortality hazards, can also both increase or decrease with age. Our results support observations that patterns of reproduction are variable across species (Fig. 2, Extended Data Fig. 1). Developing a methodology to quantify these senescence patterns of reproduction, as done here using Keyfitz’ entropy^14^ (*H*) for actuarial senescence, should be a focus of future theoretical work to unearth when, where, and what mechanisms drive how tightly the two components of senescence co-vary.

In general, our results display the discrepancy between the predictions of the classical evolutionary framework of ageing and empirical data. We suggest that researchers must widen the framework to better encompass the biology of a more diverse range of taxa. For example, the models of the classical evolutionary framework are purely age-structured, yet, in some species, demographic patterns of survival and reproduction may be influenced equally or even more by factors besides age^13^. Indeed, the force of selection does not always decline with age for some species^19^. These organisms display demographic trajectories of survival and reproduction that are better predicted by size rather than age, such as many plants^20^, which is supported by the 58% of studied plant species here that escape senescence, or corals^21^ (e.g. *Paramuricea clavata;* Fig. 1).

Many of the predictions made explicit from the classical framework of ageing have, until recently, long stood the test of time. Higher rates of extrinsic mortality, *i.e.* deaths due to the background environment, are expected to accelerate rates of senescence, whereas juvenile mortality is predicted not to play a role in the evolution of senescence^5^. Theoretical advancements, however, have shown that to have a significant effect, extrinsic mortality must be age-dependent^22^. Also, by biasing the stable age distribution of a population towards younger ages, high birth rate can also reduce the strength of selection with age^23^. The strength of selection at a given age is dependent on both the abundance of individuals in a given age class *and* the respective reproductive value of that age class^4, 23^. Following this logic, some species that display senescence yet retain high reproduction at old ages (e.g. *Marmota flaviventris* or *Pagurus longicarpus*; Fig. 1, Fig. 2) may have a stable age distribution biased towards younger individuals. This outcome would render selection too weak to promote an escape from senescence. Ultimately, how the environment shapes patterns of birth and deaths will dictate both the reproductive value of age classes and the stable age-distribution of the classes. In turn, the resulting dynamics of these pressures will affect the relative strengths of age-specific selection gradients^24^ for mortality and reproduction, and therefore patterns of senescence.

Finally, we have only considered patterns of survival and reproduction with respect to effects on the focal individual. If an individual’s survival and/or reproduction affects the fitness of others, however, and the interacting individuals are relatives, selection on the demographic age trajectories will also be weighted by these effects^25^. In our study, for example, the killer whale (*Orcinus orca*) experiences negligible actuarial senescence (*H* = 0.999) (Extended Data Fig. 1a). Killer whales are an exemplar where post-reproductive survival is hypothesised to have evolved due to the positive effects individuals can have on the survival and reproduction of offspring, *i.e.* ‘grandmother hypothesis’^4, 5, 26^. This is consistent with our results. Further evidence is also beginning to accrue elsewhere that sociality may have an important role in driving patterns of senescence beyond the remits of ‘grandmothering’^27.28^.

The emerging picture of senescence across multicellular organisms is at odds with the widely cited predictions of the classical evolutionary framework^1, 4, 5, 6^. What drives the evolution of senescence has attracted the attention of a vast research community, but we propose that the field would benefit immensely if the attention is shifted towards the underlying mechanisms allowing species to escape from senescence. We expect the greatest progress to be made by researchers honing their focus to widening the classic evolutionary theories to a framework not solely focused on age, but instead inclusive of the aforementioned factors and with a special focus on actuarial and reproductive senescence as potentially differing trajectories. Most ageing research likely stems from human desire to increase human health and life span^29^. This desire requires understanding the variation in patterns of senescence across the tree of life. For now, senescence remains an unsolved problem of biology.

## Methods

### Data

We used the COMADRE Animal Matrix Database^2^ (v. 3.0.0) and COMPADRE Plant Matrix Database^3^ (v. 5.0.0) to obtain age trajectories of survival and reproduction. These open-access data repositories consist of matrix population models^13^ (MPMs) incorporating high-resolution demographic information on the survival and reproduction patterns of over 1,000 animal and plant species worldwide^2, 3^. Both databases include information on species for which the data have been digitised and thoroughly error-checked. In addition, we contacted authors for clarifications when any doubt about the interpretation of the life cycle of the species emerged. We imposed a series of selection criteria to restrict our analyses to data of the highest quality possible.

i. MPMs were parameterised with field data from non-disturbed, unmanipulated populations (*i.e.* natural populations) to best describe the species’ age trajectories.
ii. MPMs had dimension ≥3 × 3 (i.e. rows × columns). Generally, low dimensions MPMs lack quality for the estimation of life history traits^30^. This selection criterion also helps avoid problems with quick convergence to stationary equilibrium, at which point the estimates of life history trait values and rates of senescence become unreliable^8, 31^.
iii. MPMs were only used when the entire life cycle was explicitly modelled including recordings of survival, development, and reproduction for all life cycle stages.
iv. When MPMs existed for multiple populations within a given species, we calculated the arithmetic element-by-element mean MPM to obtain a single MPM per species.
v. When multiple studies existed for the same species, we considered only the study of greater duration to ensure the highest temporal variation in the population dynamics was captured.
vi. Studies of annual plant species modelled using seasonal projection matrices were not included; we chose only species using an annual time step. This is due to the difficulties of converting their population dynamics to an annual basis to compare with all other species’ models.
vii. Included MPMs have stage-specific survival values ≤ 1. In a small number of ≤ published models, the stage-specific survival values can exceed 1 due to clonality being hidden in the matrix, rounding errors, or other mistakes in the original model^2, 3^.
viii. MPMs were from species of which phylogenetic data was available, to ensure we were able to account for phylogenetic relatedness on our models.

The result of these criteria was a subset of 308 species of animals and plants from the initial databases, which we used for our analysis. Of these, 48 were animals, with 13 invertebrates and 35 vertebrates. The remaining 260 species were plants, with 23 gymnosperms and 237 angiosperms. We provide a list of all the species used, their categorisation as displaying or escaping senescence including value of *H*, and their relevant source study (Supplementary Table 1).

### Displaying *vs* escaping senescence

MPMs are a summary of the population dynamics of a given species, from which we can calculate several life history traits. To do so, we first must decompose an MPM (***A)*** into its sub-components^13^:

***U*** – containing the stage-specific survival rates

***F*** – containing the stage-specific per-capita reproduction rates

***C*** – containing stage-specific per-capita clonality rates

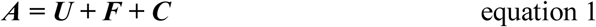

This decomposition facilitates the estimation of key life history traits, including Keyfitz’ entropy (*H*)^14^. Calculating *H* requires first obtaining the age-specific survivorship curve l(x)from ***U***. First, the definition of age requires a choice of a stage that corresponds to “birth”. Following Jones *et al*.^8^, we defined the stage corresponding to birth as the first established non-propagule stage (e.g., not seeds or seed bank in the case of plants, nor larvae or propagules in animals) due to the estimate uncertainty of parameters involved in those stages. The calculation of l(x) was then implemented according to Caswell (p. 118-21)^13^.

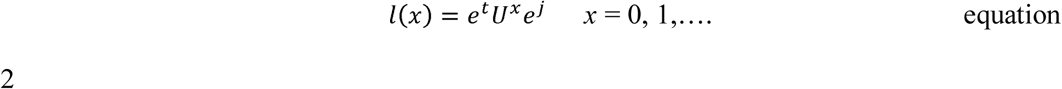

We considered survivorship trajectories beginning at the age of maturity (calculated following 5.47–5.54 in Caswell^13^) and ending at the age at which 5% survivorship from maturity occurs. This is because a cohort modelled by iteration of the ***U*** matrix eventually decays exponentially at a rate given by the dominant eigenvalue of ***U***, and converges to a quasi-stationary distribution given by the corresponding right eigenvector **w**. Once this convergence has happened, mortality remains constant with age, and so to prevent our conclusions being overly influenced by this assumption, we calculated the age at which the cohort had converged to within a specified percentage (5%) of the quasi-stationary distribution^8, 31^.

*H* is proportional to the area under the age-specific survivorship curve and is calculated as follows

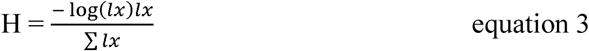

If most mortality occurs later in life, *H* <1, and individuals in the population display actuarial senescence. On the contrary, if *H*>1, the risk of mortality declines with age and the individuals in the population escape actuarial senescence. Here, we categorise species as either escaping or displaying senescence, however, values of H∼1 are more likely to indicate negligible senescence, where risk of mortality remains relatively constant with age.

### Phylogenetic analyses for actuarial senescence

After categorising species as displaying (*H*<1) or escaping (*H*>1) senescence, we accounted for the phylogenetic relatedness of the species studied to determine the influence of a species’ evolutionary history in its likelihood of evolving or escaping senescence. To build the phylogenetic tree with the species included in this study, we used data from the Open Tree of Life^32^ (OTL), a database that combines public available taxonomic and phylogenetic information across the tree of life. We first checked that the species names in our data were taxonomically accepted using the *taxize* R package^33^. Then, we obtained animal and plant phylogenetic trees form OTL for the list of species in our data using *rotl* R package^34^. The branch length of the resulting tree was computed using the *compute.brlen* function from the R package *ape*^35^, with Grafen’s^36^ arbitrary branch lengths. Polytomies (*i.e.* >2 species with the same ancestor) were resolved using the function multi2di from *ape* package^35^, which transforms polytomies into a series of random dichotomies with one or several branches of length zero.

To evaluate the role of phylogenetic relatedness in determining the patterns of variation of actuarial senescence we estimated Pagel’s λ^15^. This metric is an index bounded between zero and one, where values ∼0 indicate that the evolutionary history of the species explains little about the variation of the trait measured, and values ∼1 suggest that that evolutionary history mostly explains the observed variation of the trait across the studied species. To estimate Pagel’s λ we used the R package *motmot.2.0*^37^. A full summary of the phylogenetic signals obtained for each of the four monophyletic groups can be found in the Extended Data and Figures (Table 1).

### Reproduction analysis

We calculated reproductive age-trajectories for the species in our analysis to investigate whether reproductive senescence followed the same pattern as actuarial senescence in species that display *vs.* escape actuarial senescence. Age-specific reproduction (*m(x)*) was calculated following Caswell (p. 118-21)^13^. Briefly, the proportional structure of the cohort at age *x* is given by

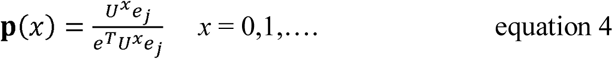

The total sexual reproductive output per individual at age *x* is given by

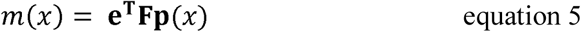

For two species, *Araucaria mulleri* and *Juniperus procera, m(x)* was incalculable, and so both species’ *l(x)* and *m(x)* trajectories are not reported. For the remaining 306 species, the *l(x)* and *m(x)* trajectories are found in the Extended Fig. 1a-h.

## Code availability

Code used for analysis is available in the supplementary information.

## Data availability

Data are available from the source data files in the supplementary information and from the COMPADRE Plant Matrix Database and COMADRE Animal Matrix Database (www.compadre-db.com).

## Acknowledgements

M.R was supported by funding from the Biotechnology and Biological Sciences Research Council (BBSRC) [grant number BB/M011224/1]. P.C. was supported by a Ramón Areces Foundation Postdoctoral Scholarship. This research emerged through funding by NERC R/142195-11-1 to R.S-G.

## Author information

### Contributions

M.R. and R.S.G. conceived the project. M.R. and P.C conducted the analysis and produced all visualizations. M.R drafted the first version and together with P.C. and R.S.-G., revised and edited the manuscript.

## Extended data figures and tables

**Fig. 1:**
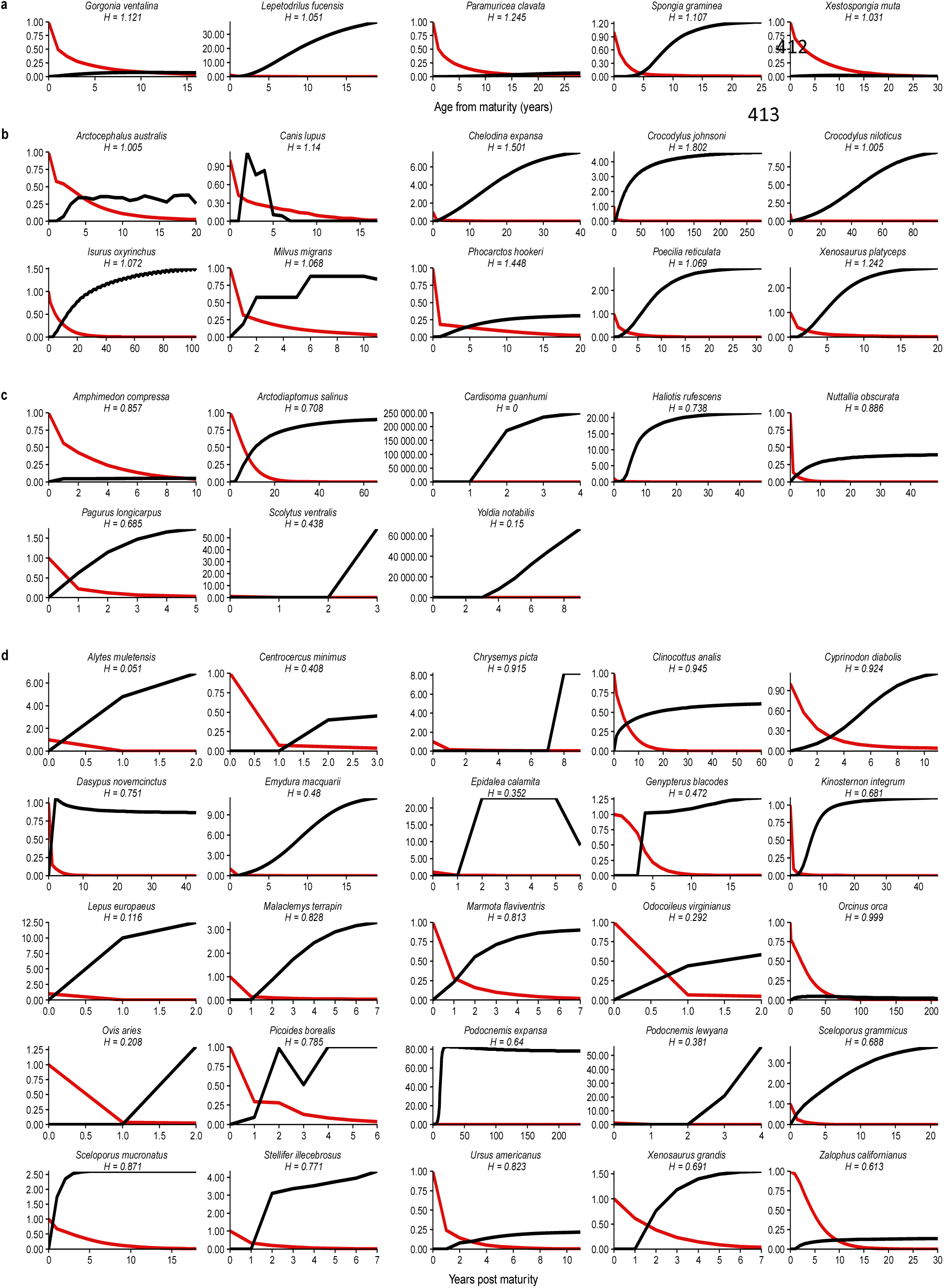
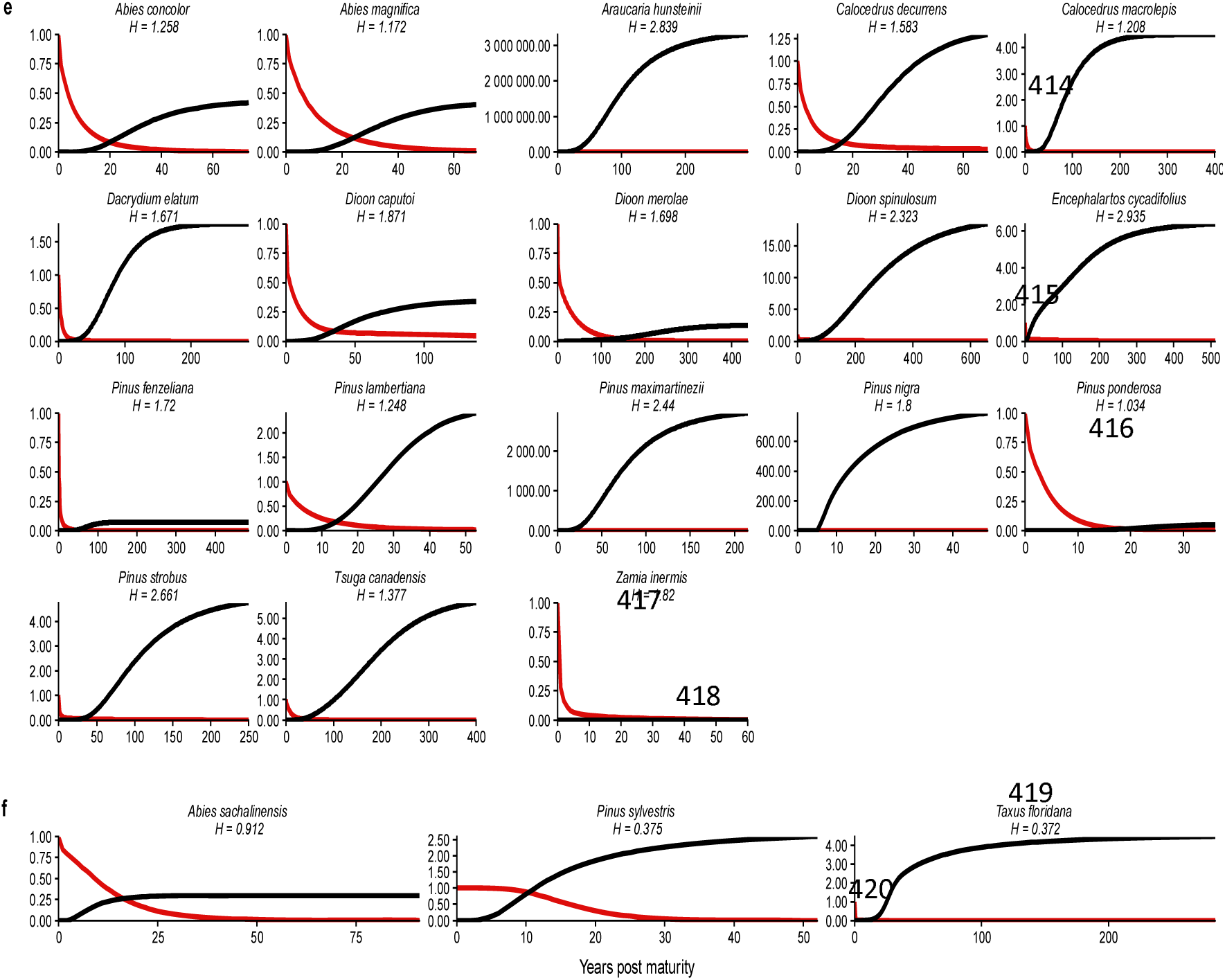
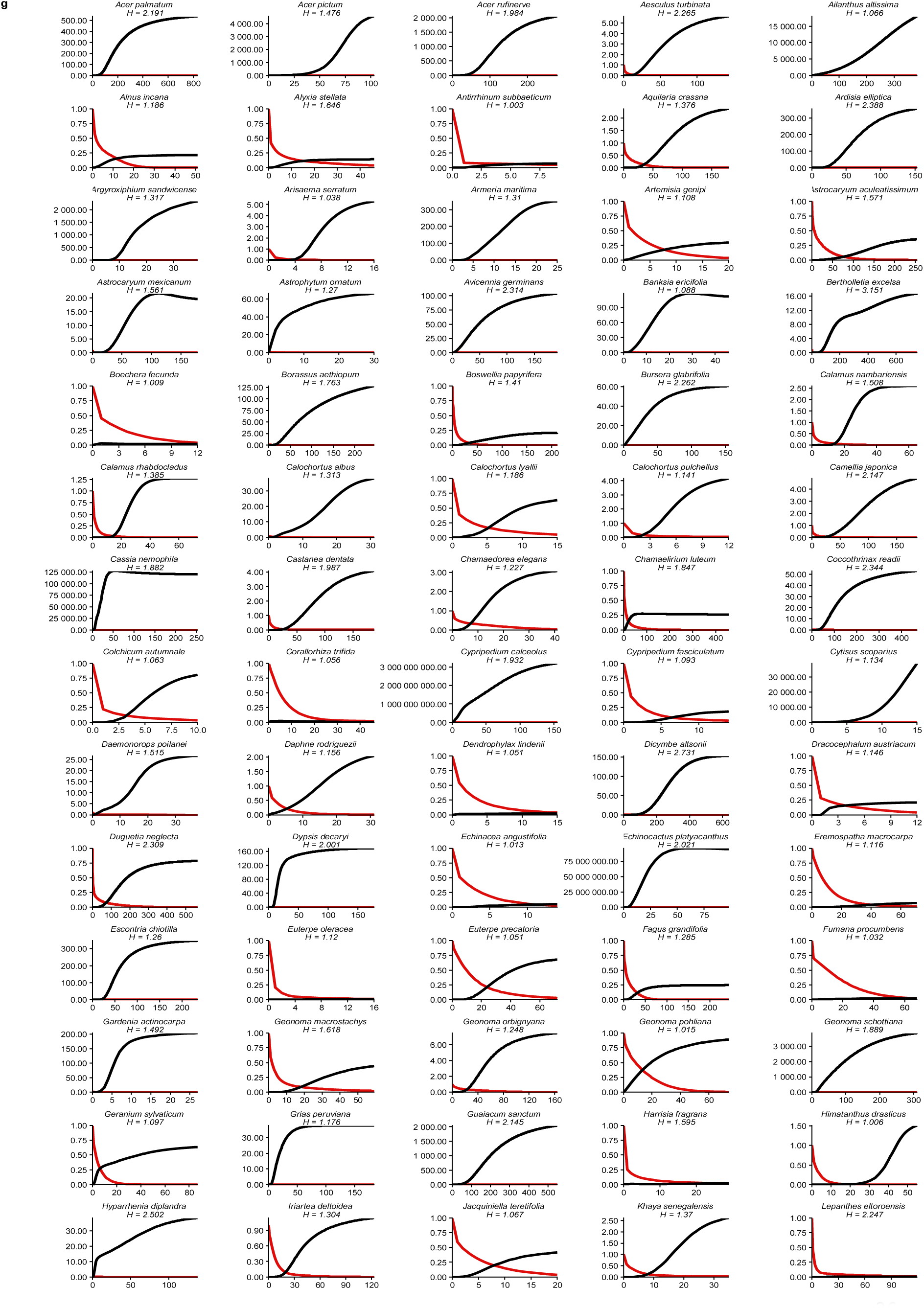
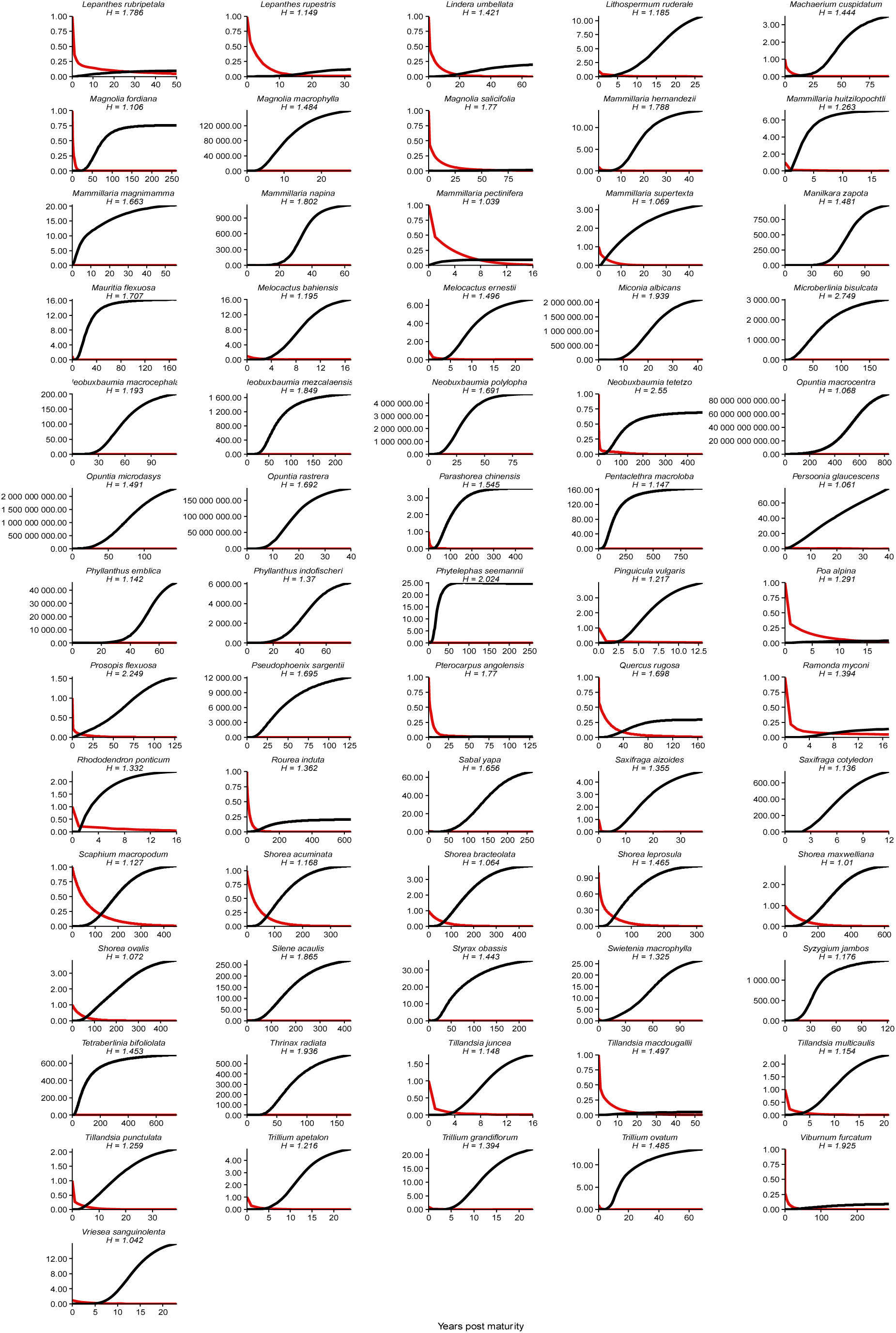
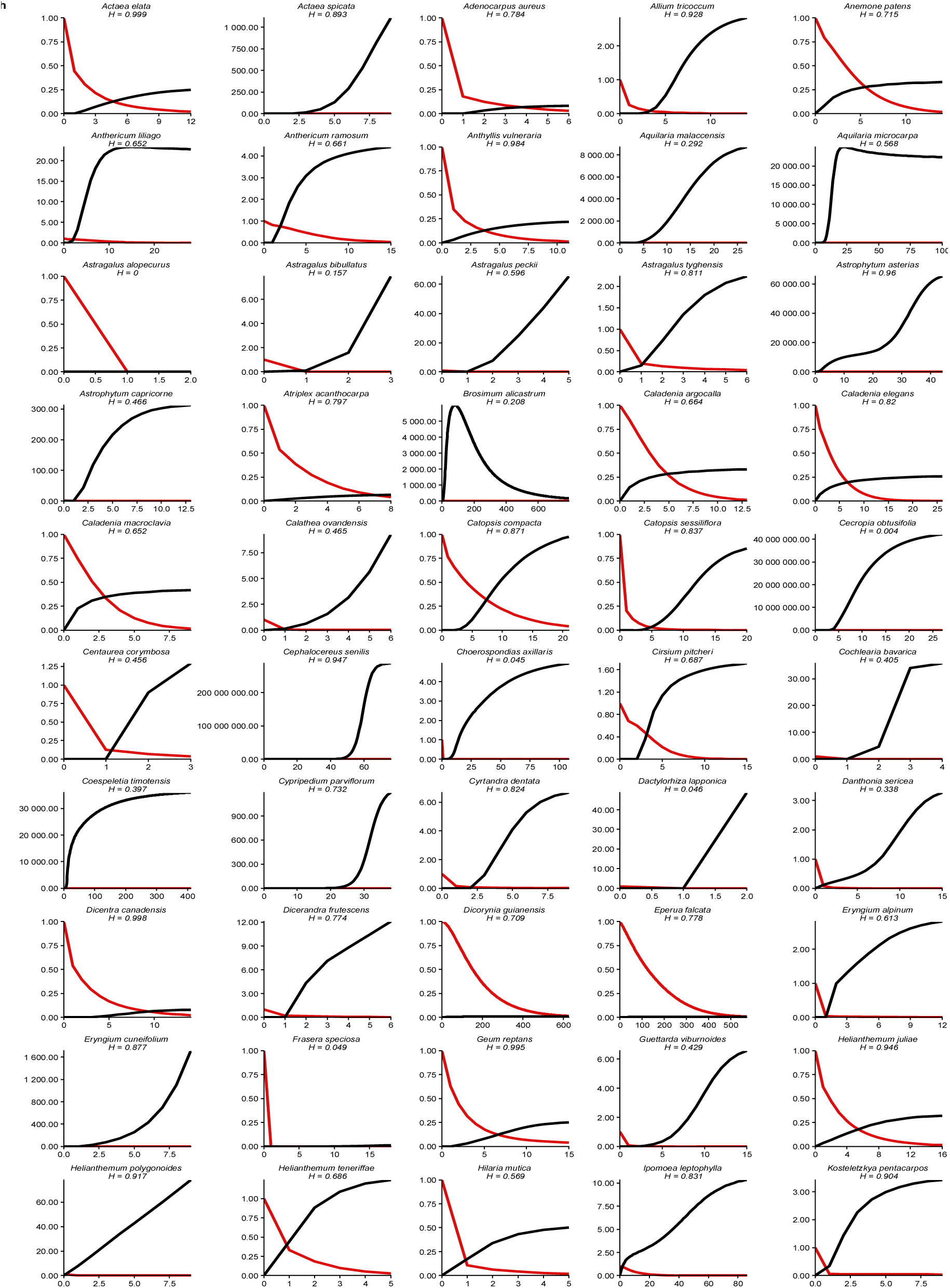
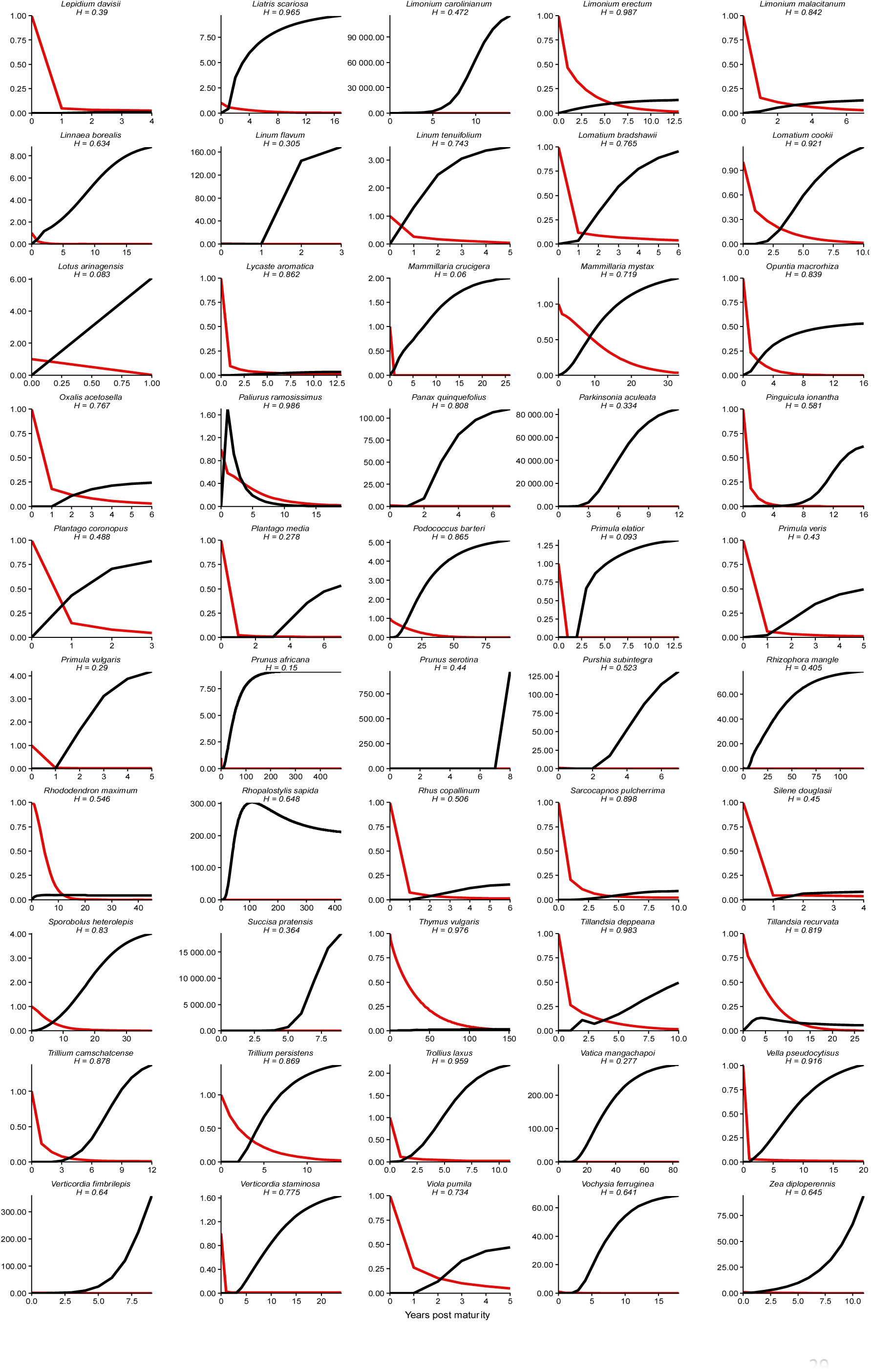
Survivorship (red) and reproduction (black) age-trajectories for 306 of our study species. **a)** Invertebrates that escape senescence **b)** Vertebrates that escape senescence **c)** Invertebrates that display senescence **d)** Vertebrates that escape senescence **e)** Gymnosperms that escape senescence **f)** Gymnosperms that display senescence **g)** Angiosperms that escape senescence **h)** Angiosperms that display senescence. Trajectories are conditional upon reaching, and are shown from, the age of maturity, labelled as 0, to the age at which 5% of the mature cohort is still alive. The mature cohort is defined to have survivorship of 1.

**Extended Data Table 1:**
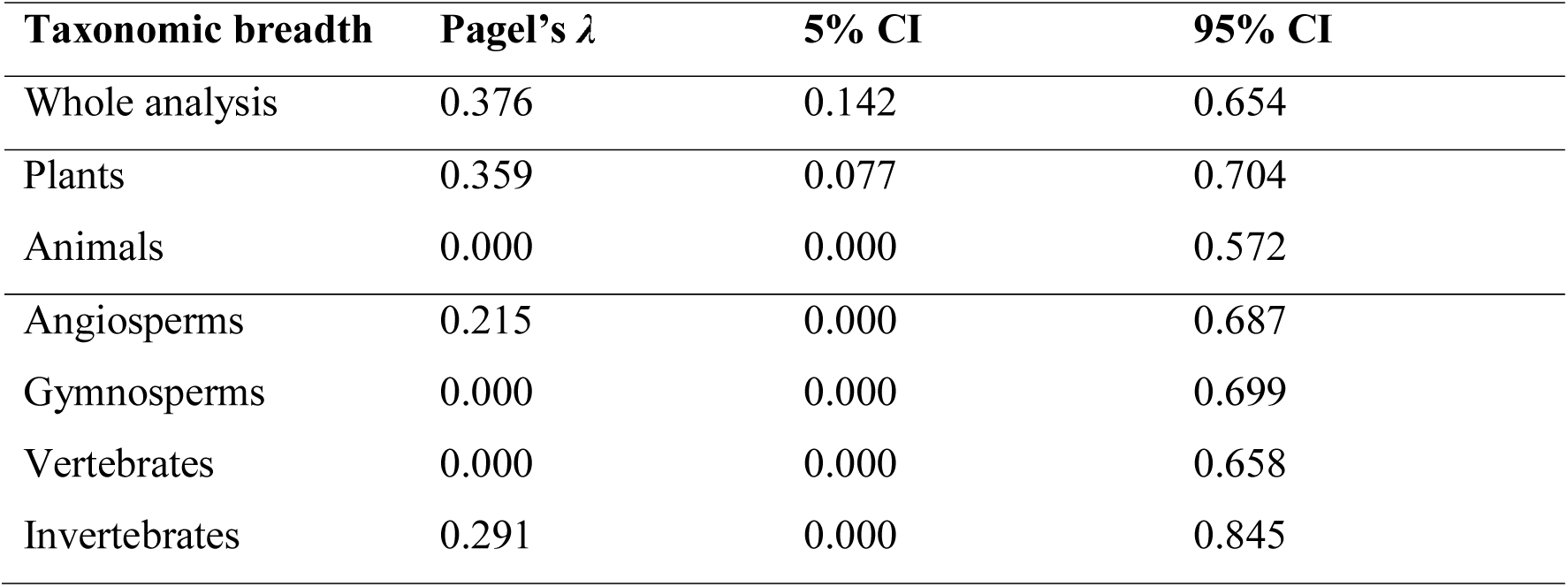
Phylogenetic signals of actuarial senescence for each major taxonomic group of our 308 studied species are relatively weak. Calculated estimates of Pagel’s λ within the phylogenetic analysis of actuarial senescence across our study species. Pagel’s λ is an index bounded between zero and one, where values ∼0 indicate that the evolutionary history of the species explains little about the variation of the trait measured, and values ∼1 suggest that that evolutionary history mostly explains the observed variation of the trait across the studied species.

## Supplementary information

**Supplementary Table 1:**
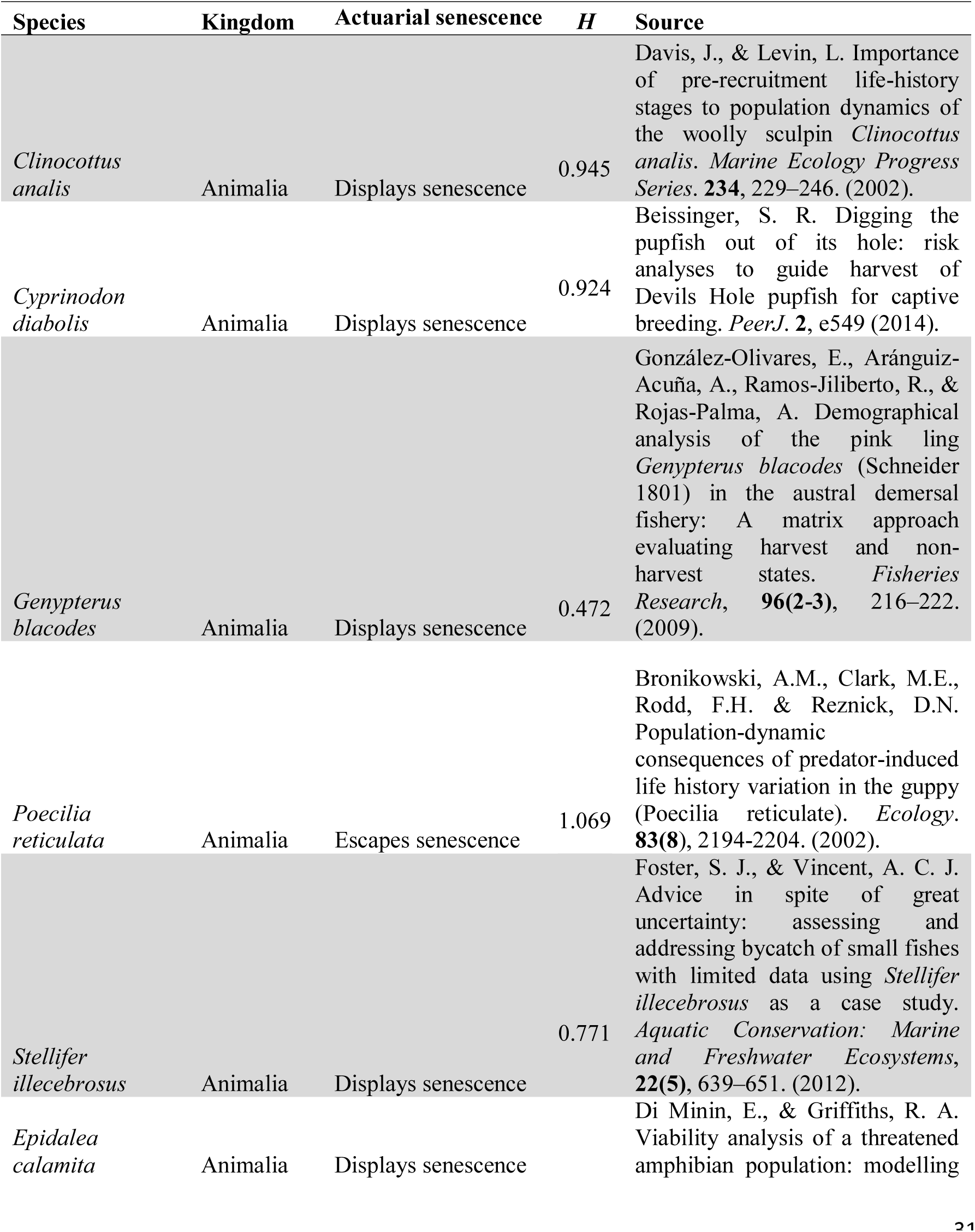
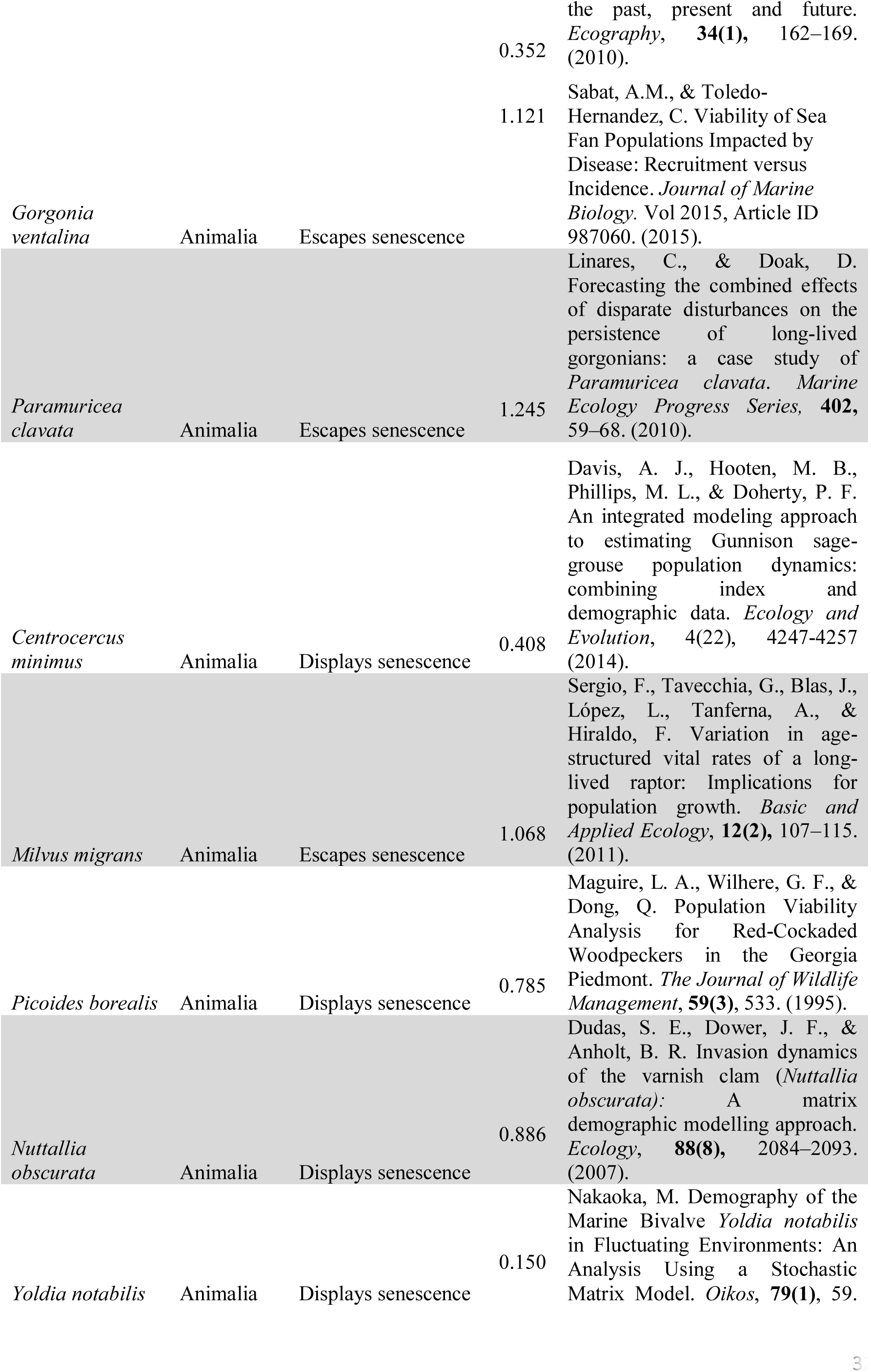
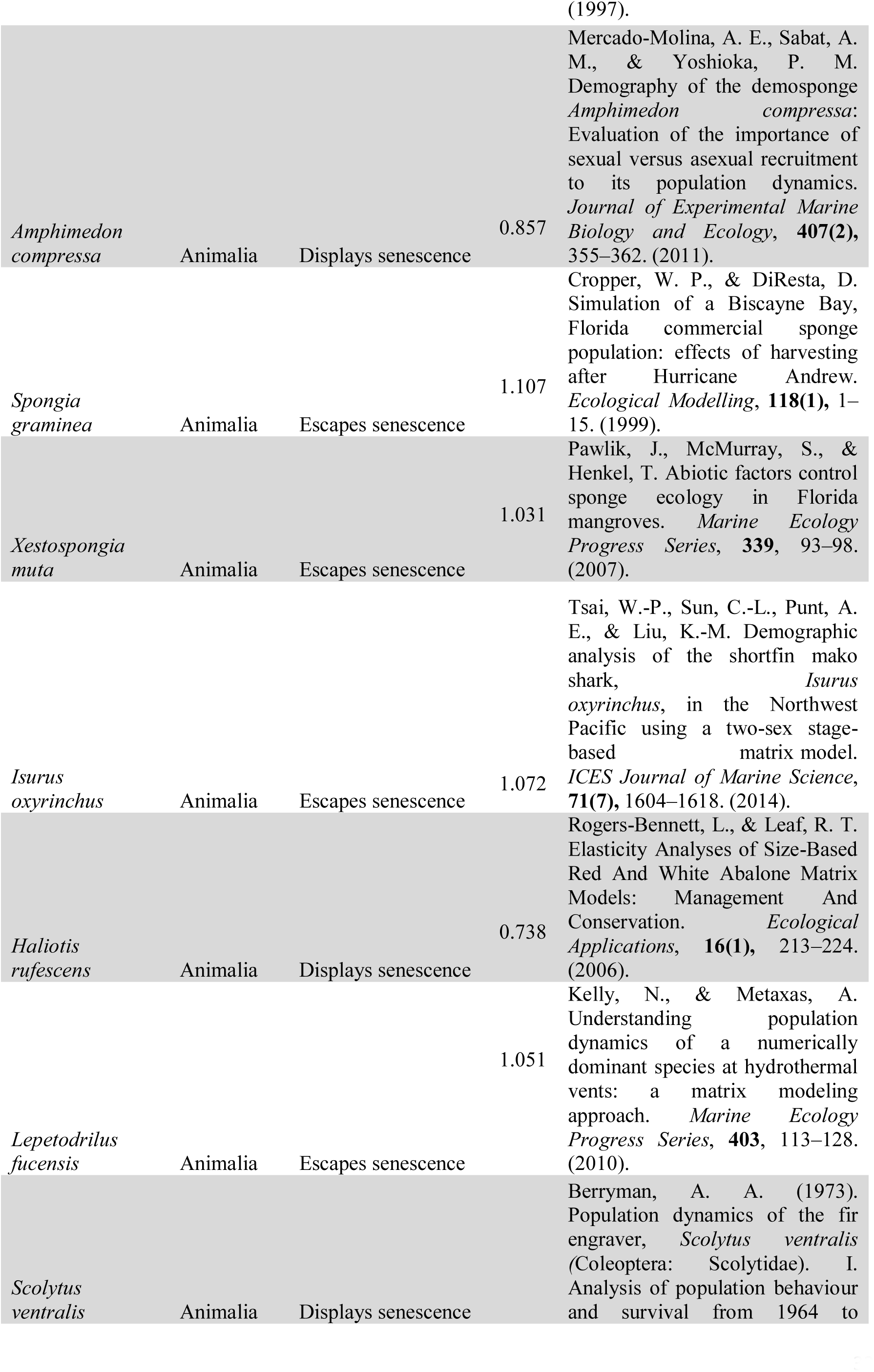
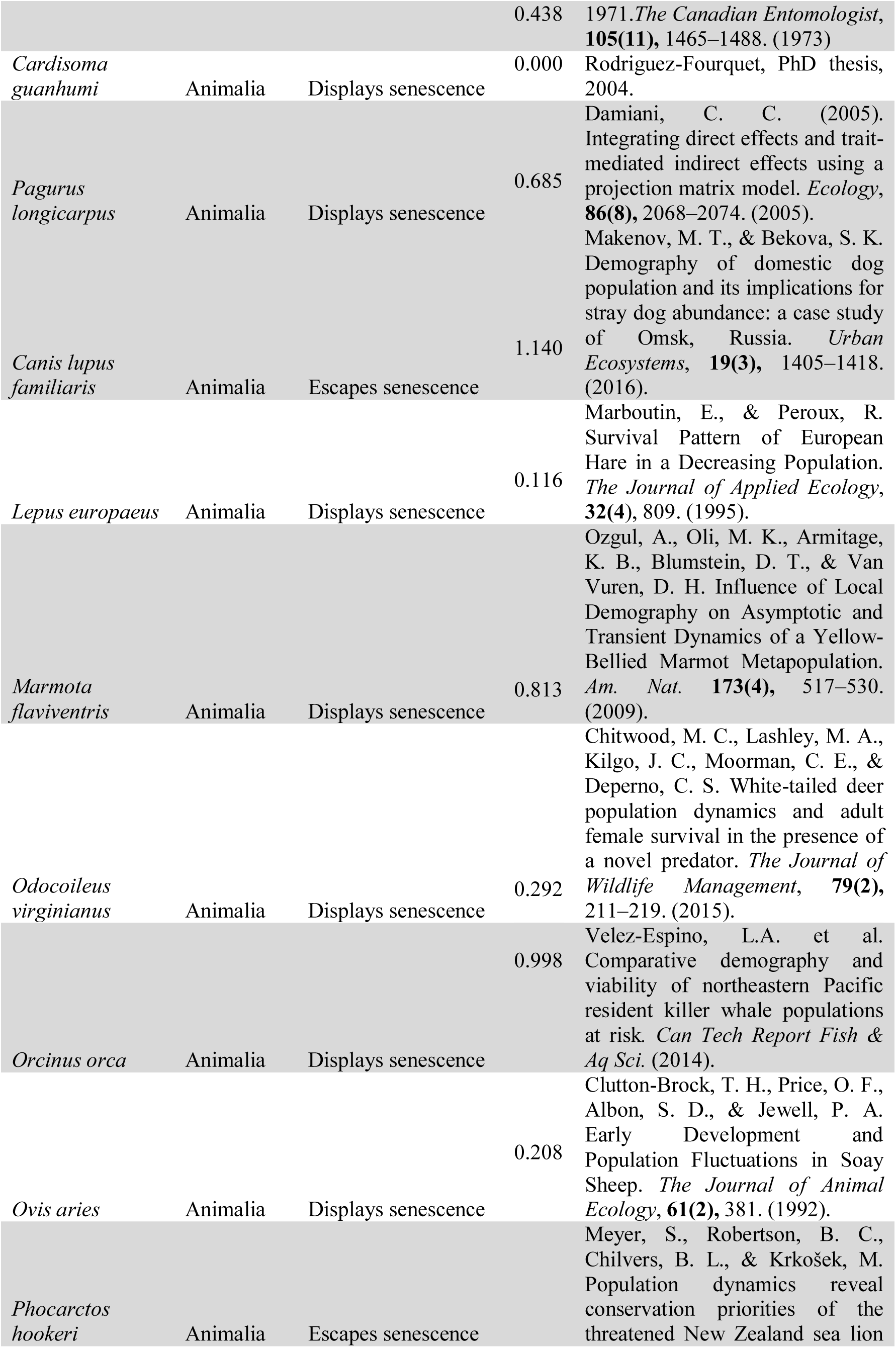
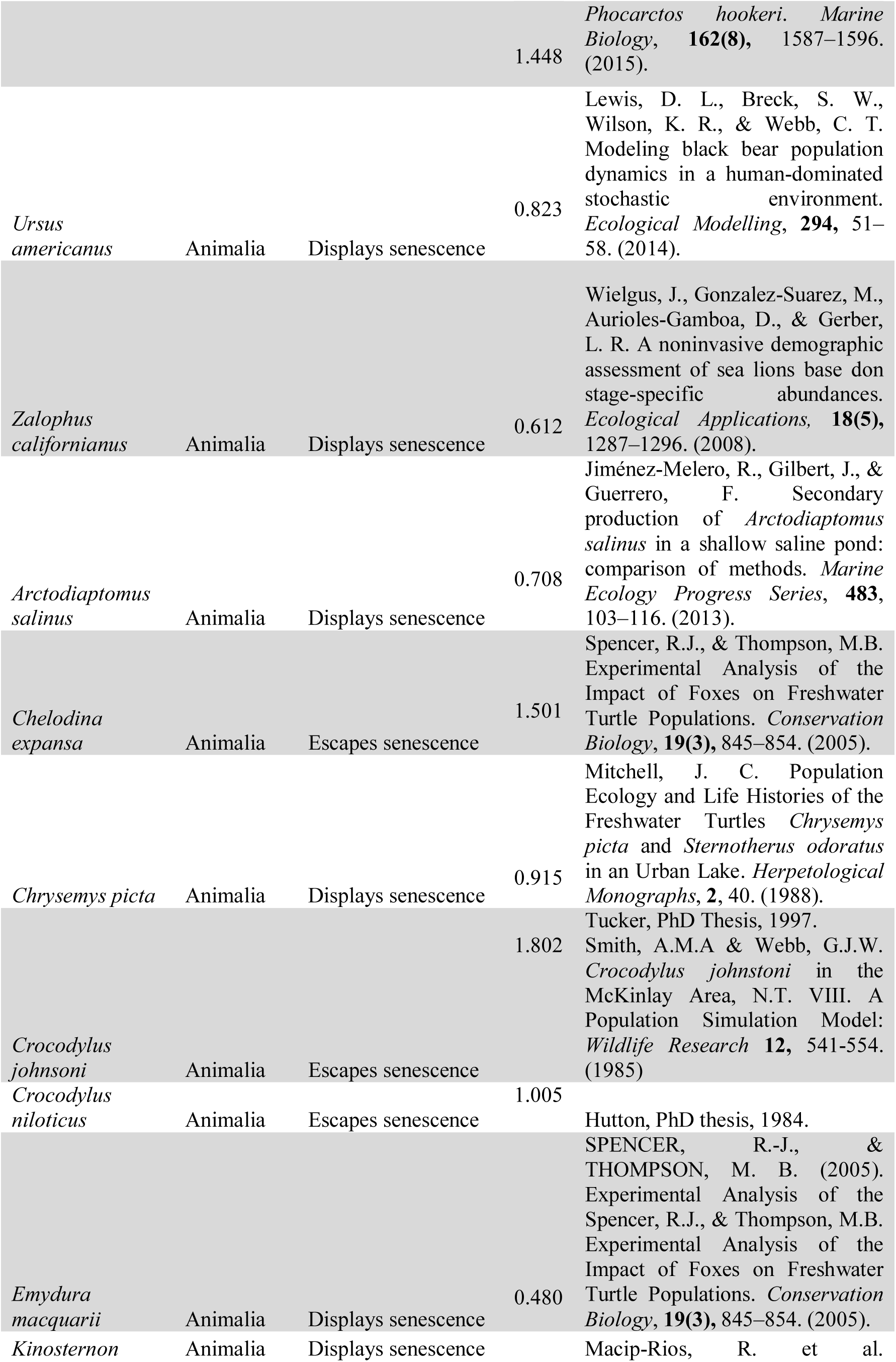
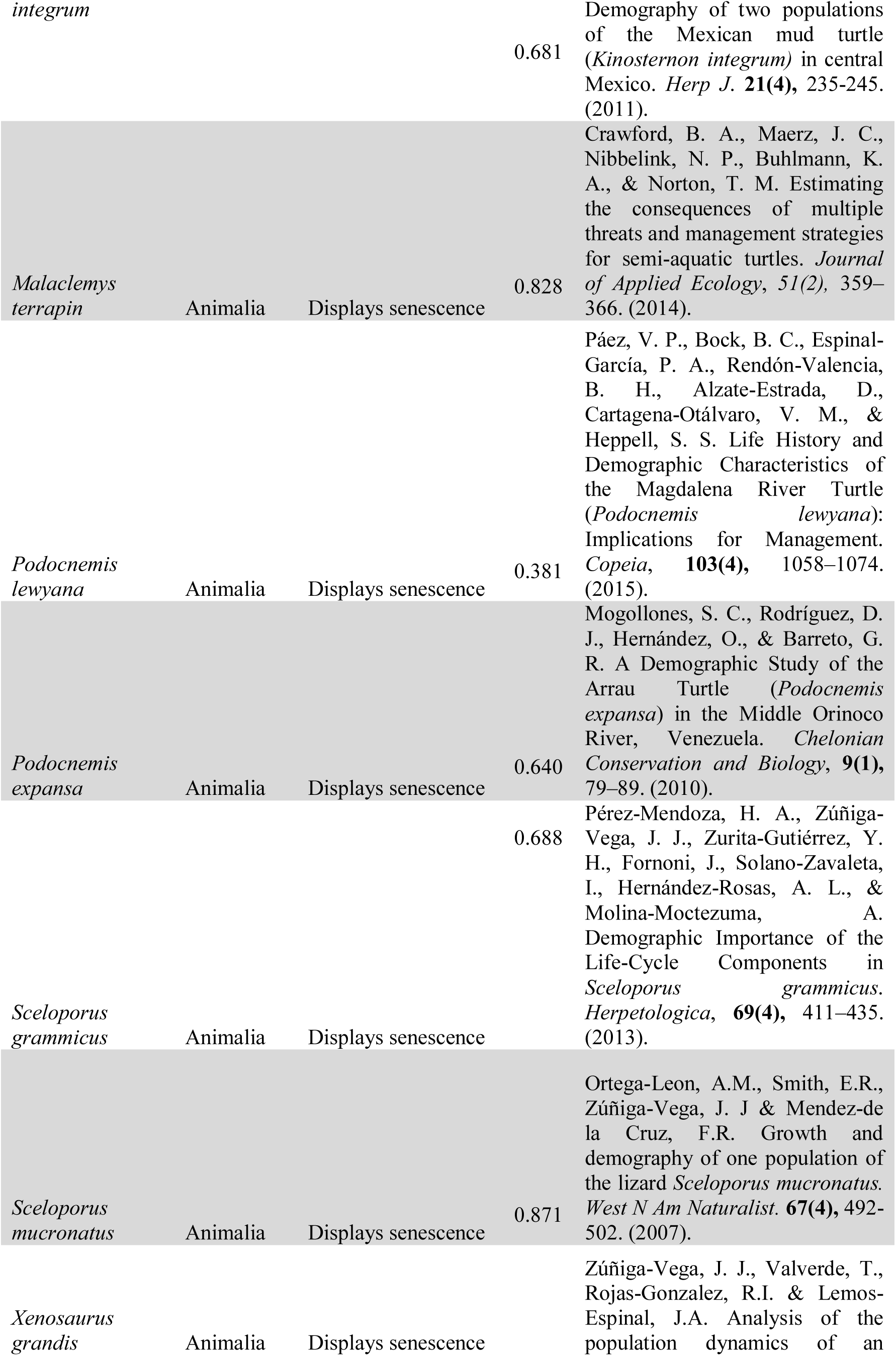
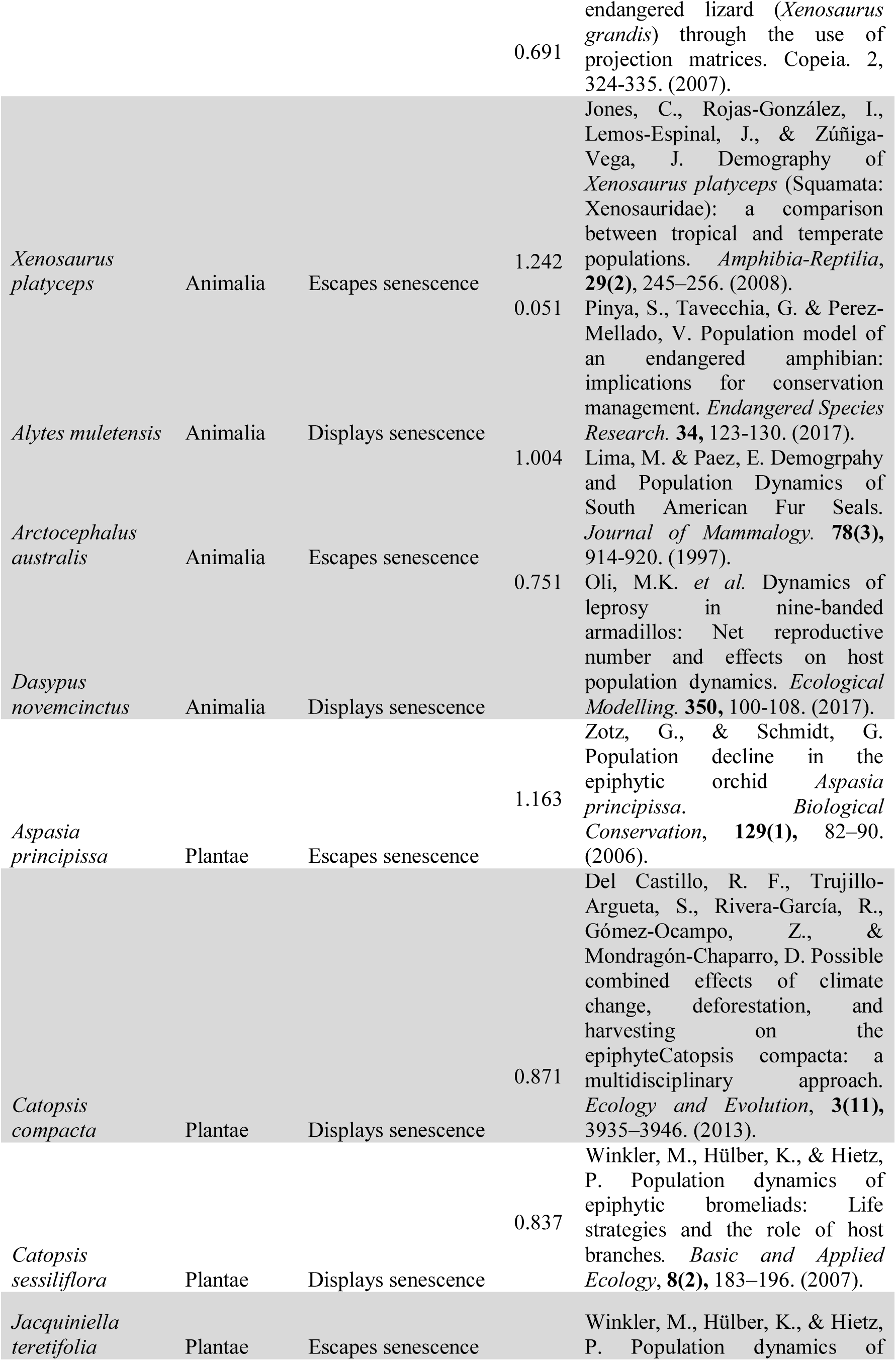
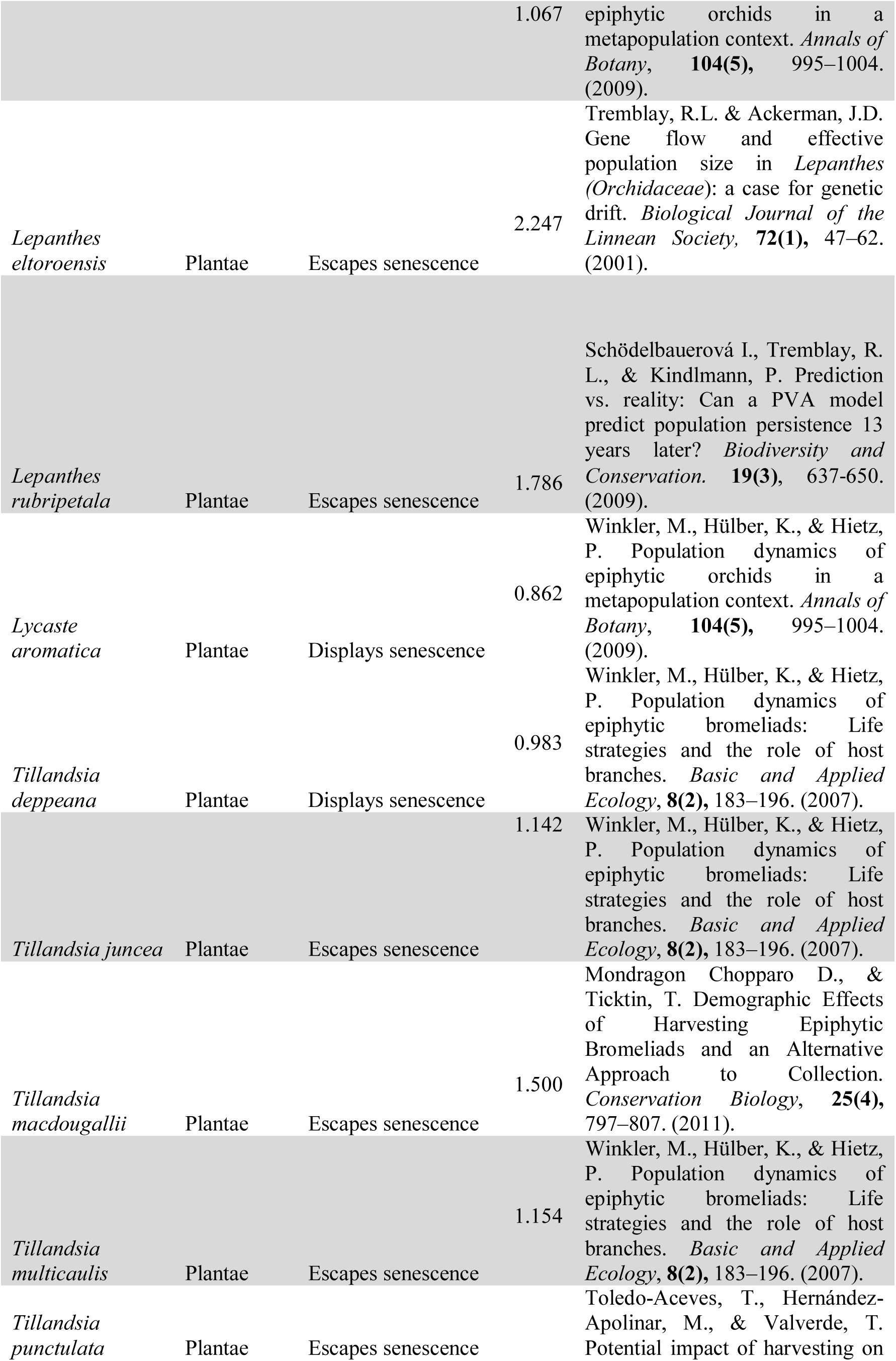
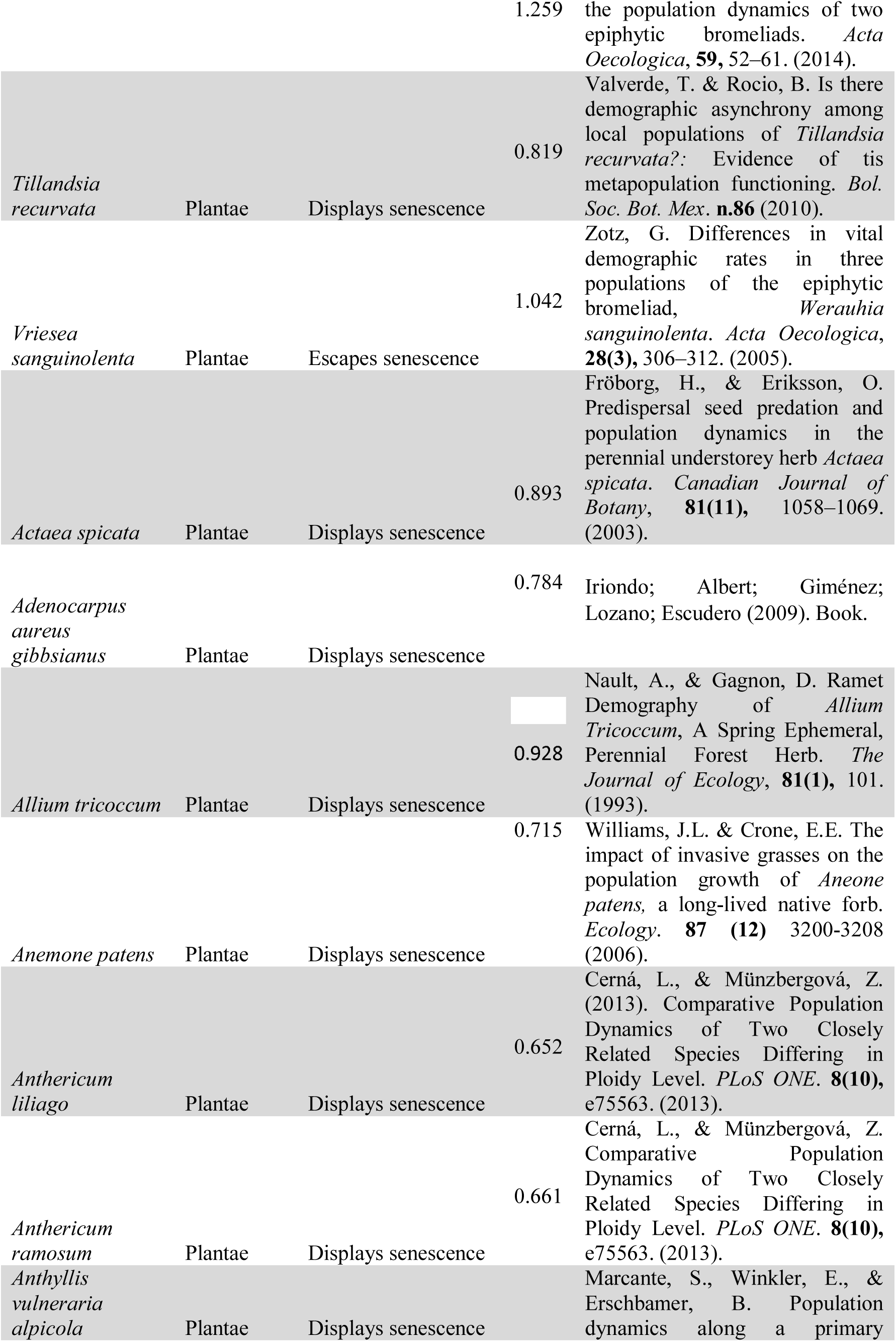
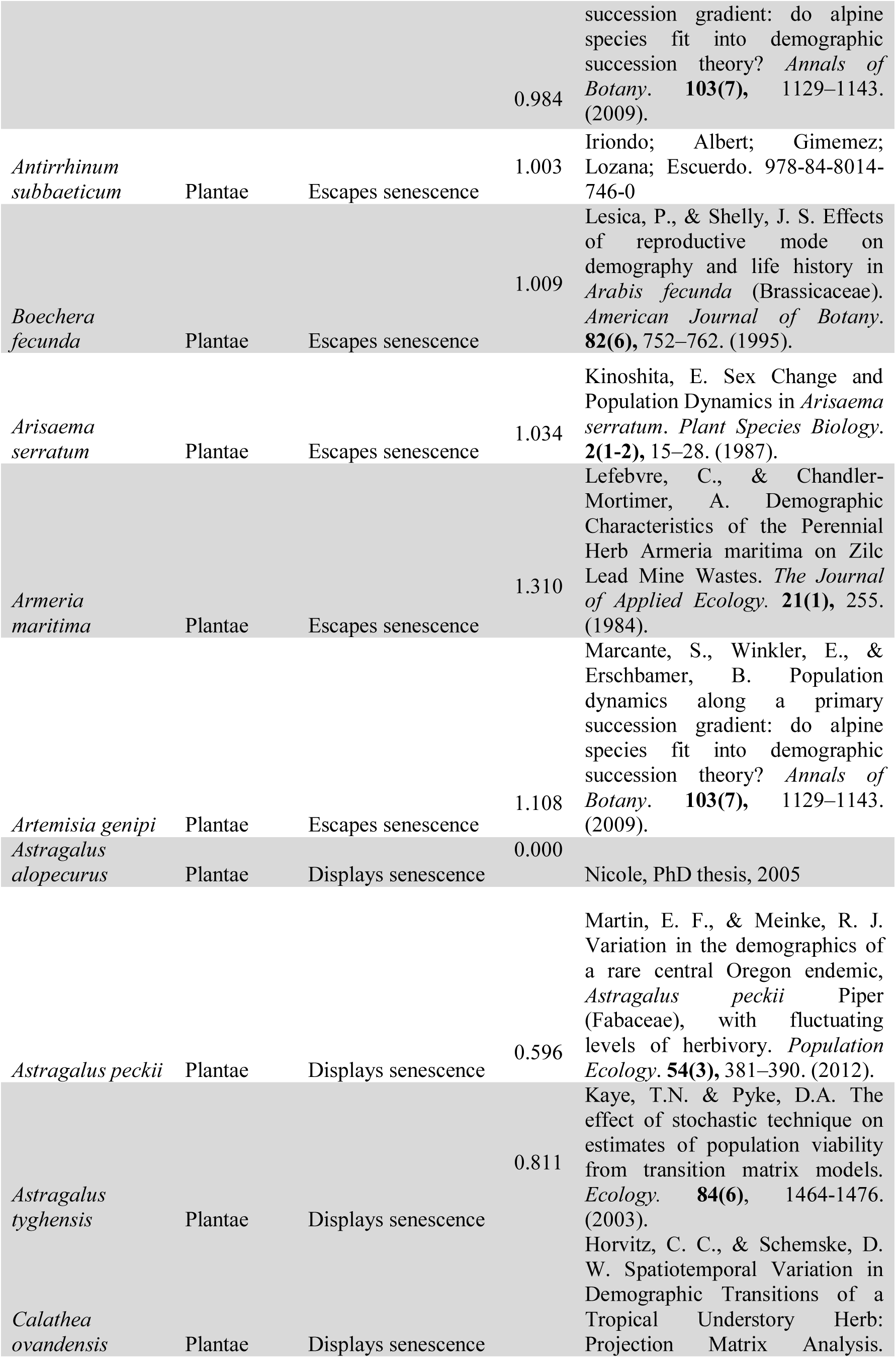
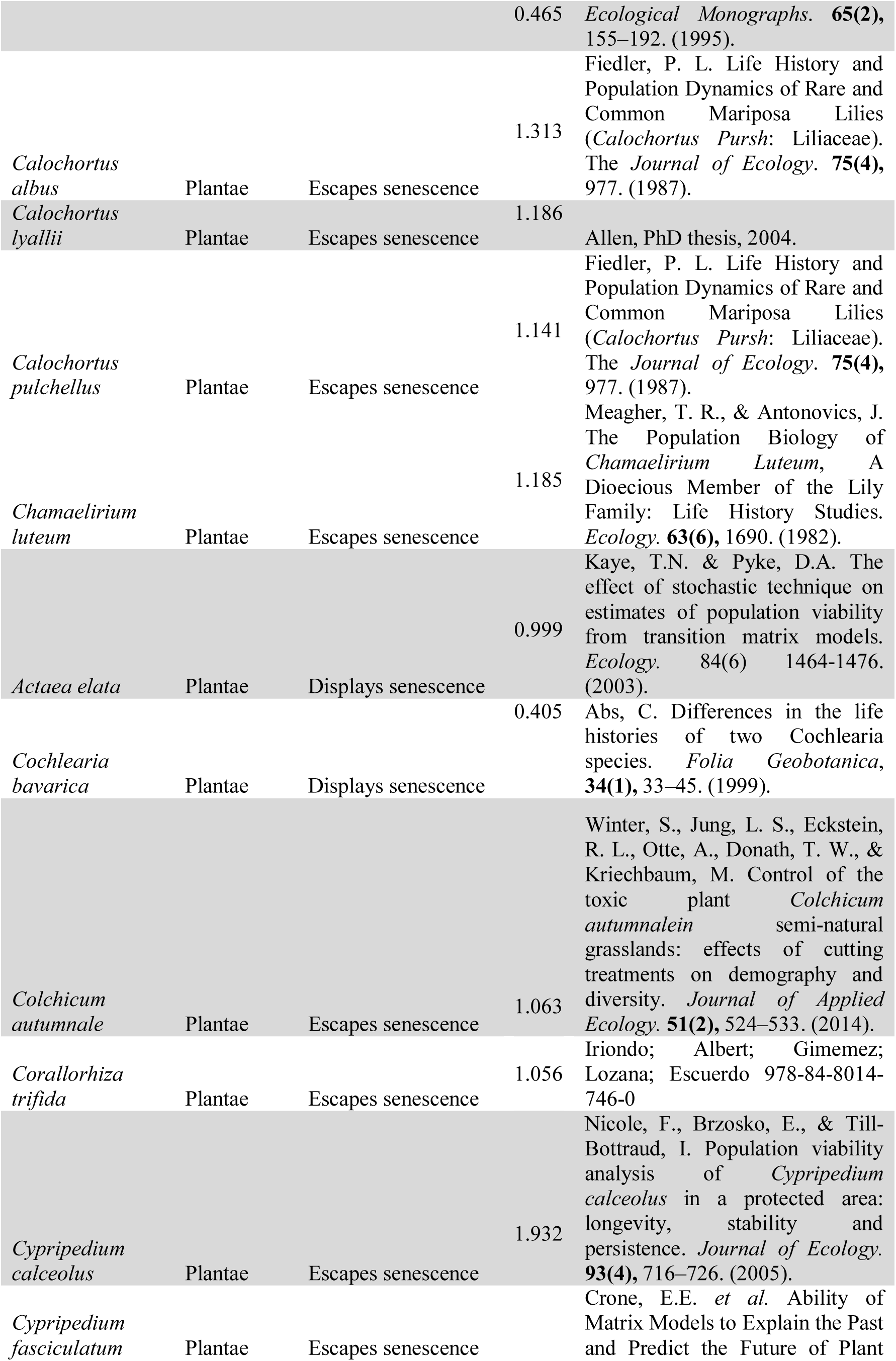
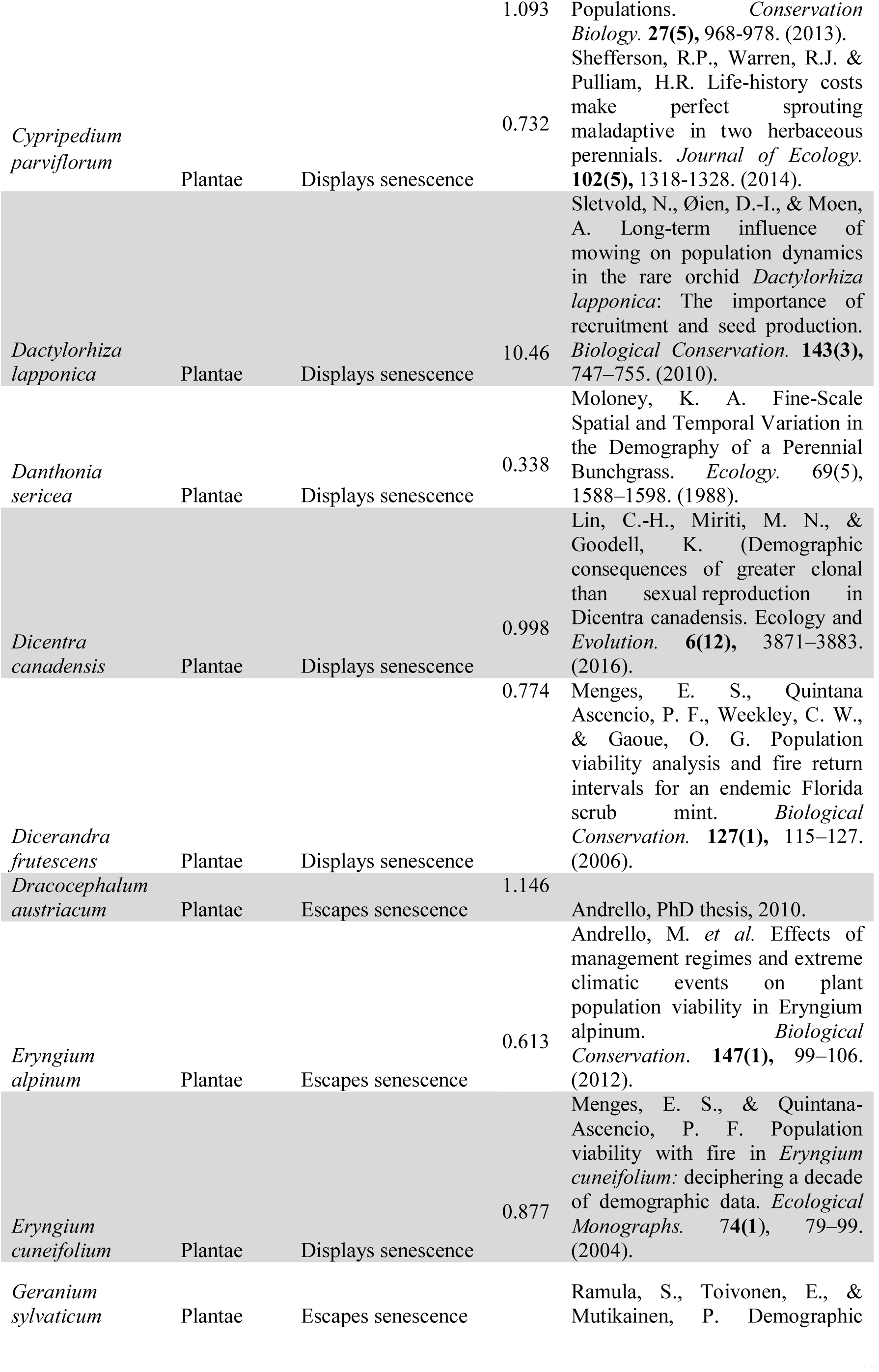
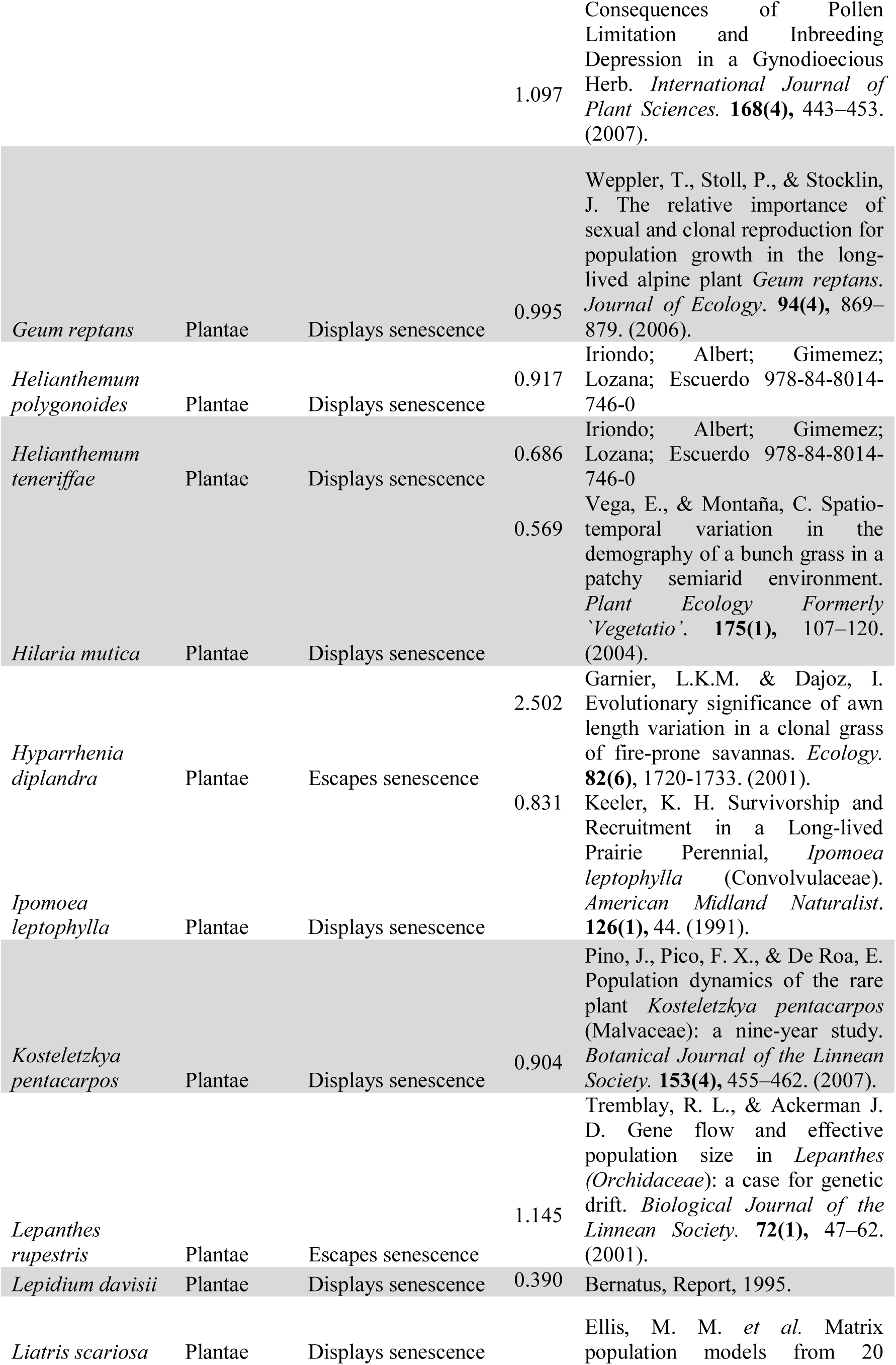
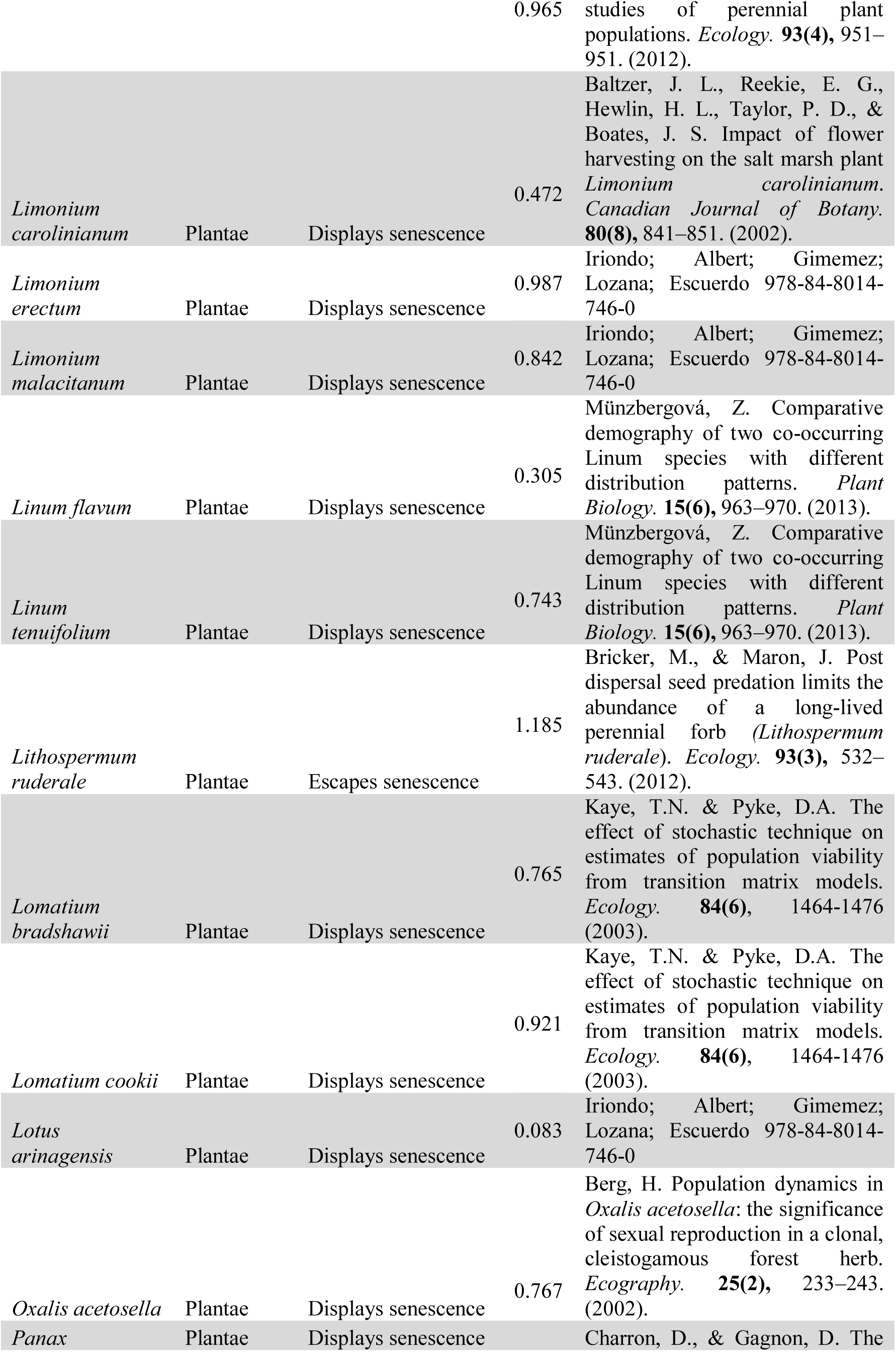
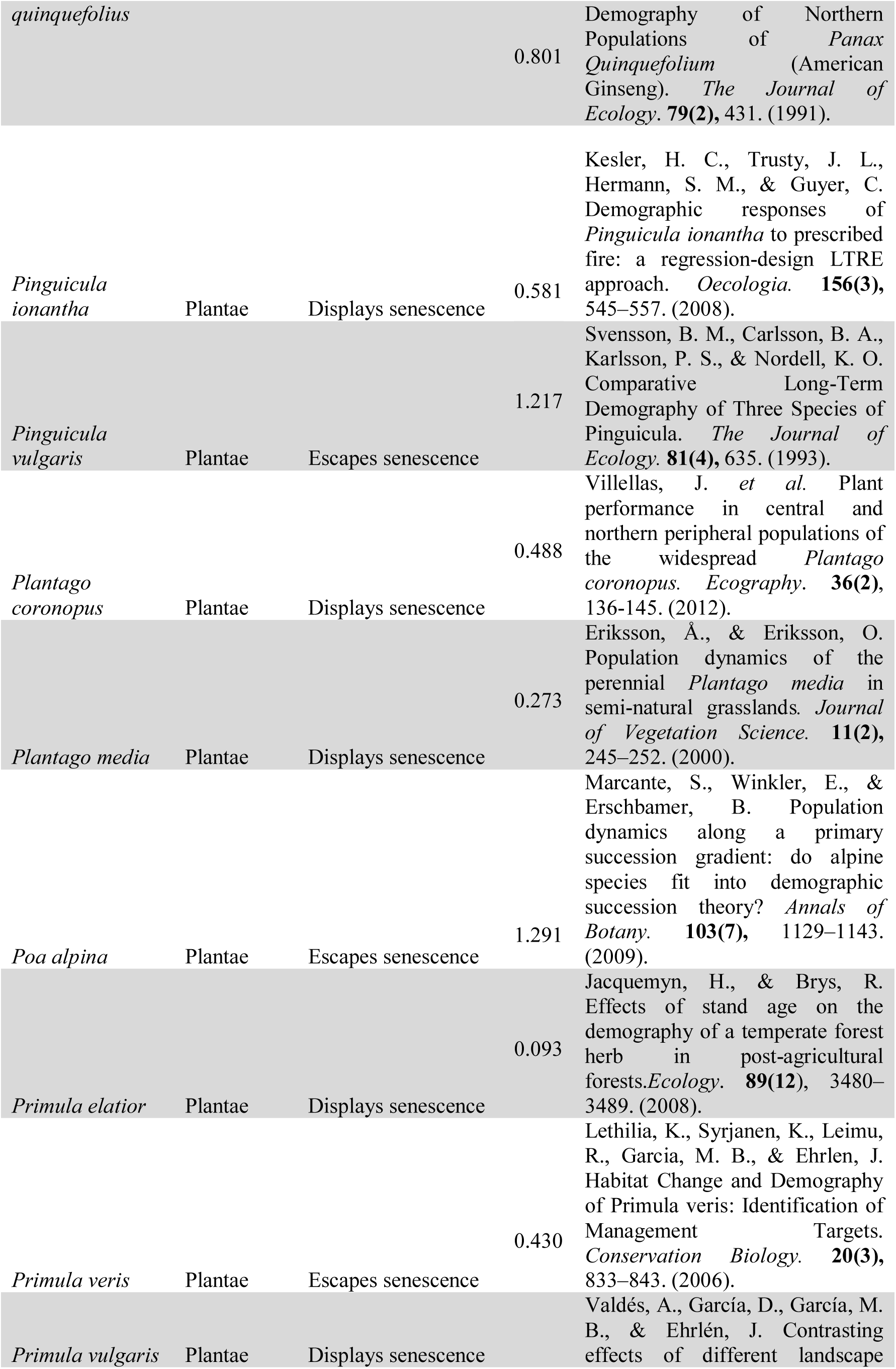
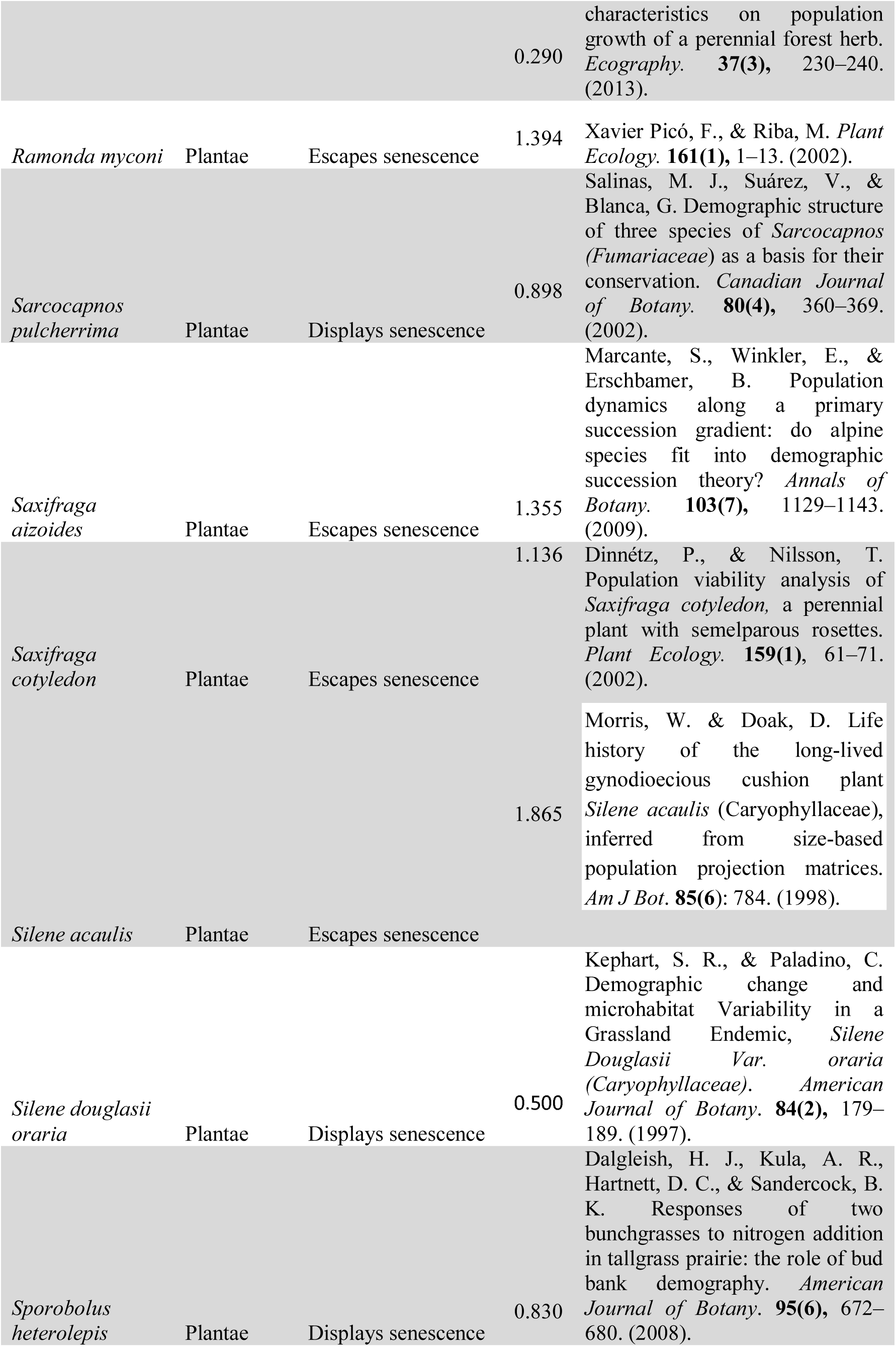
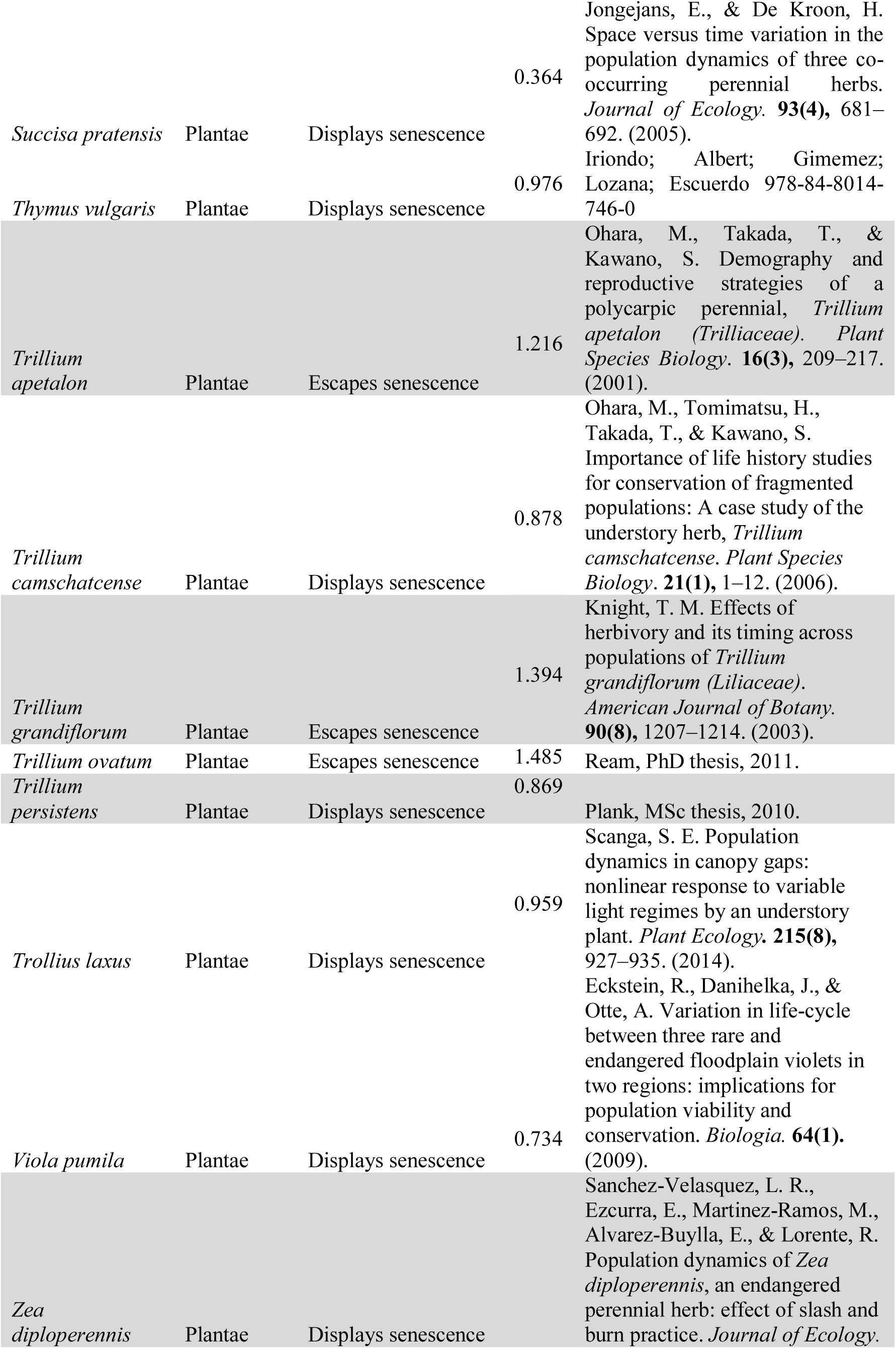
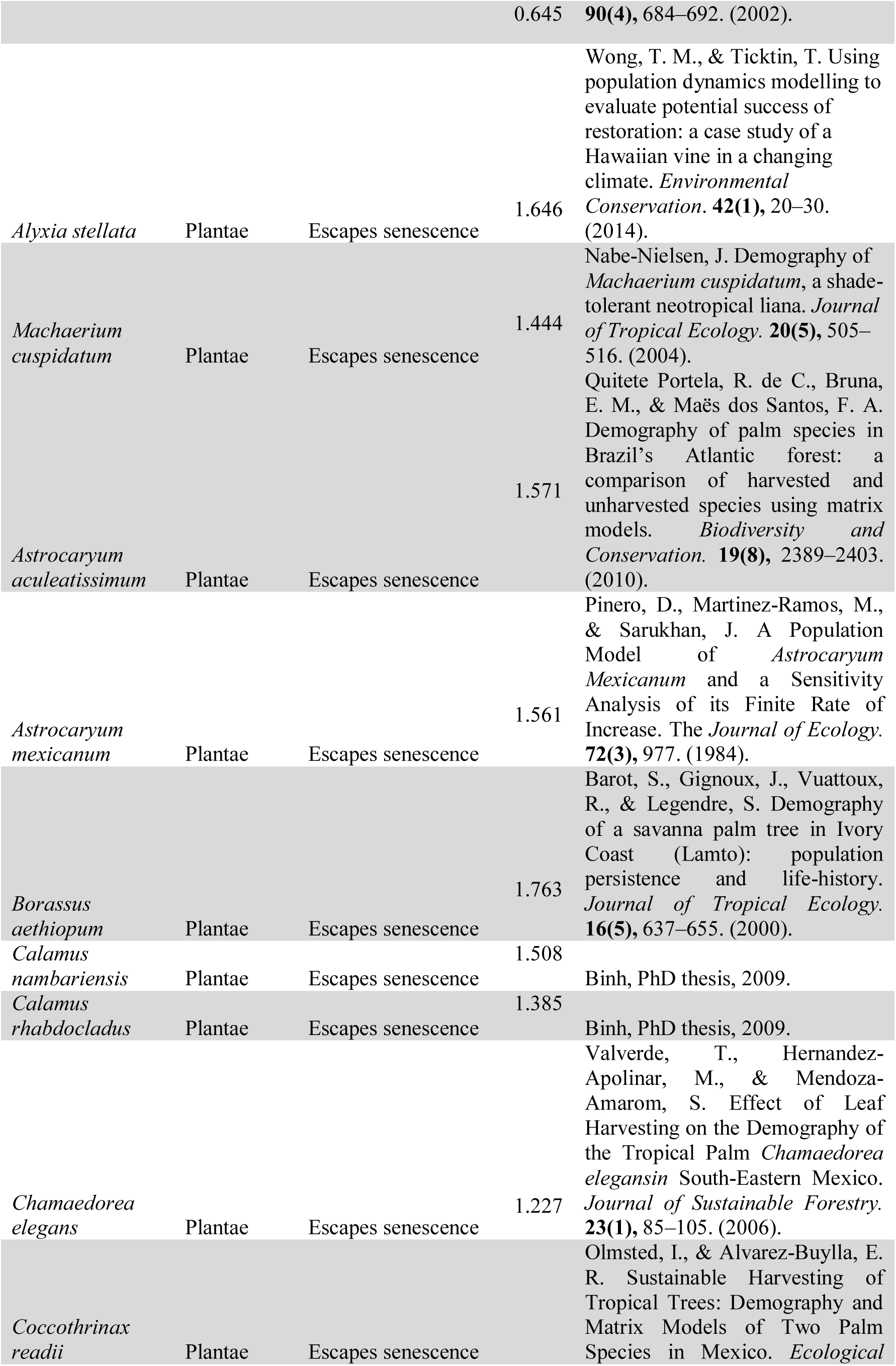
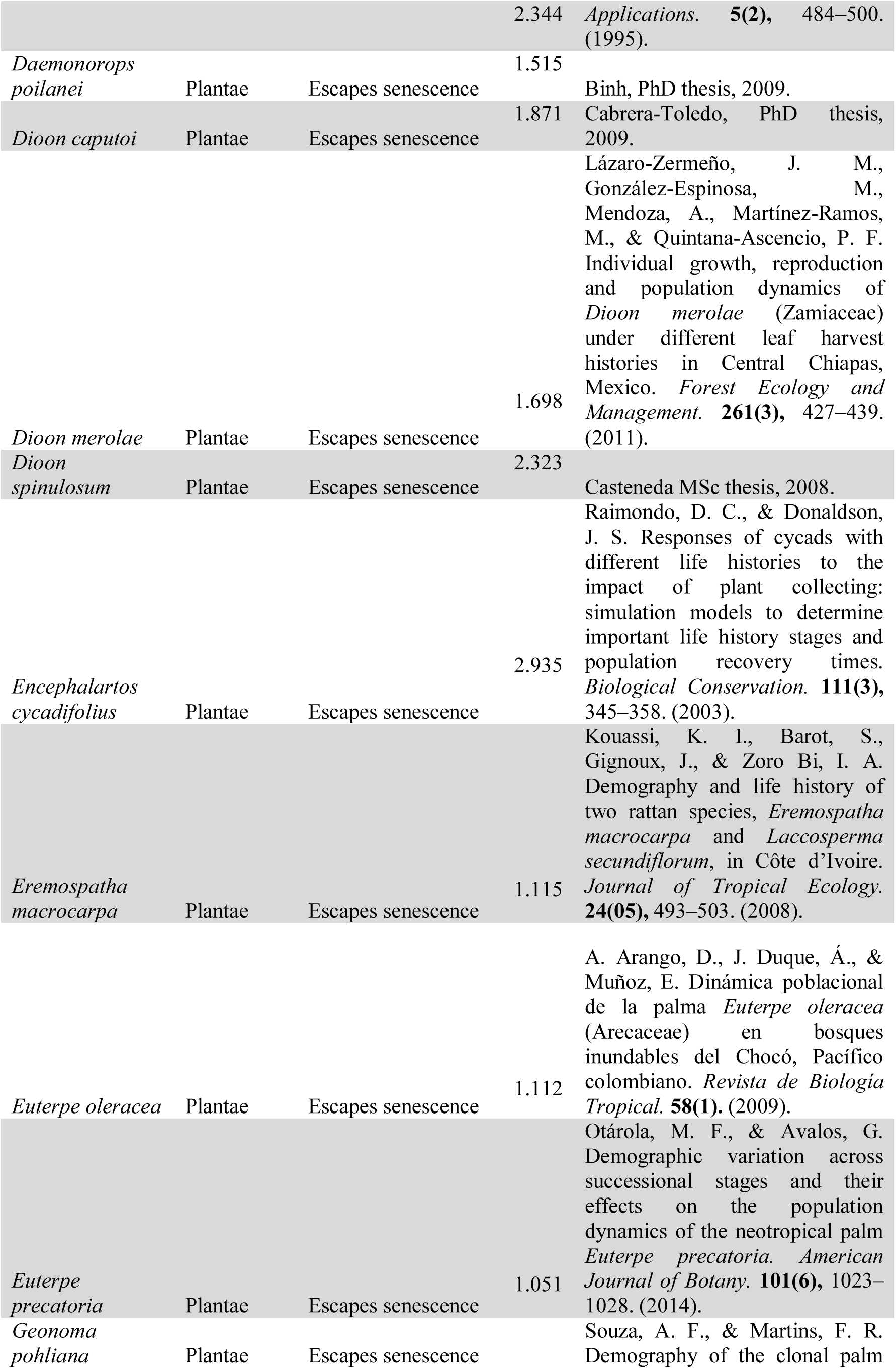
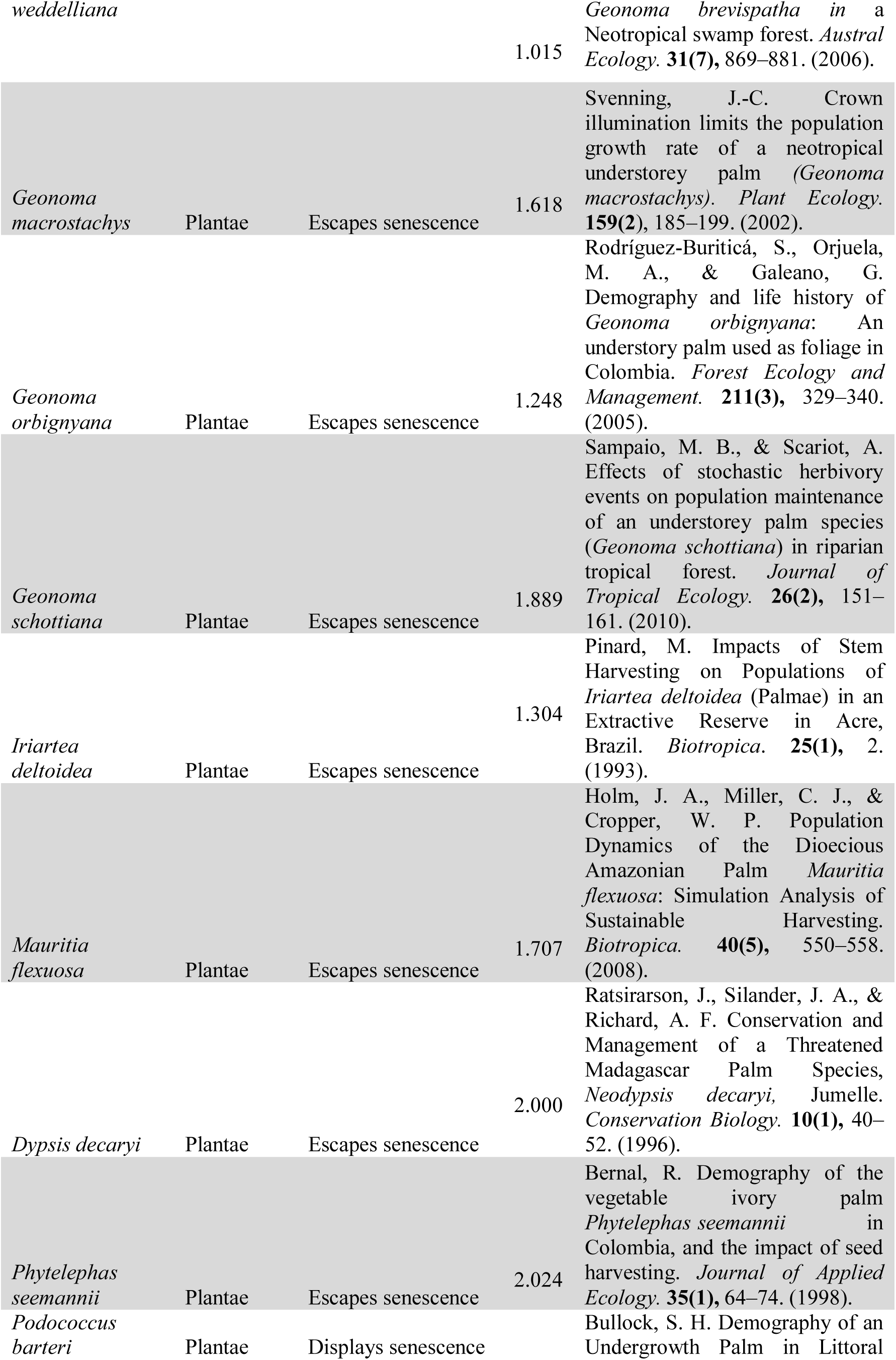
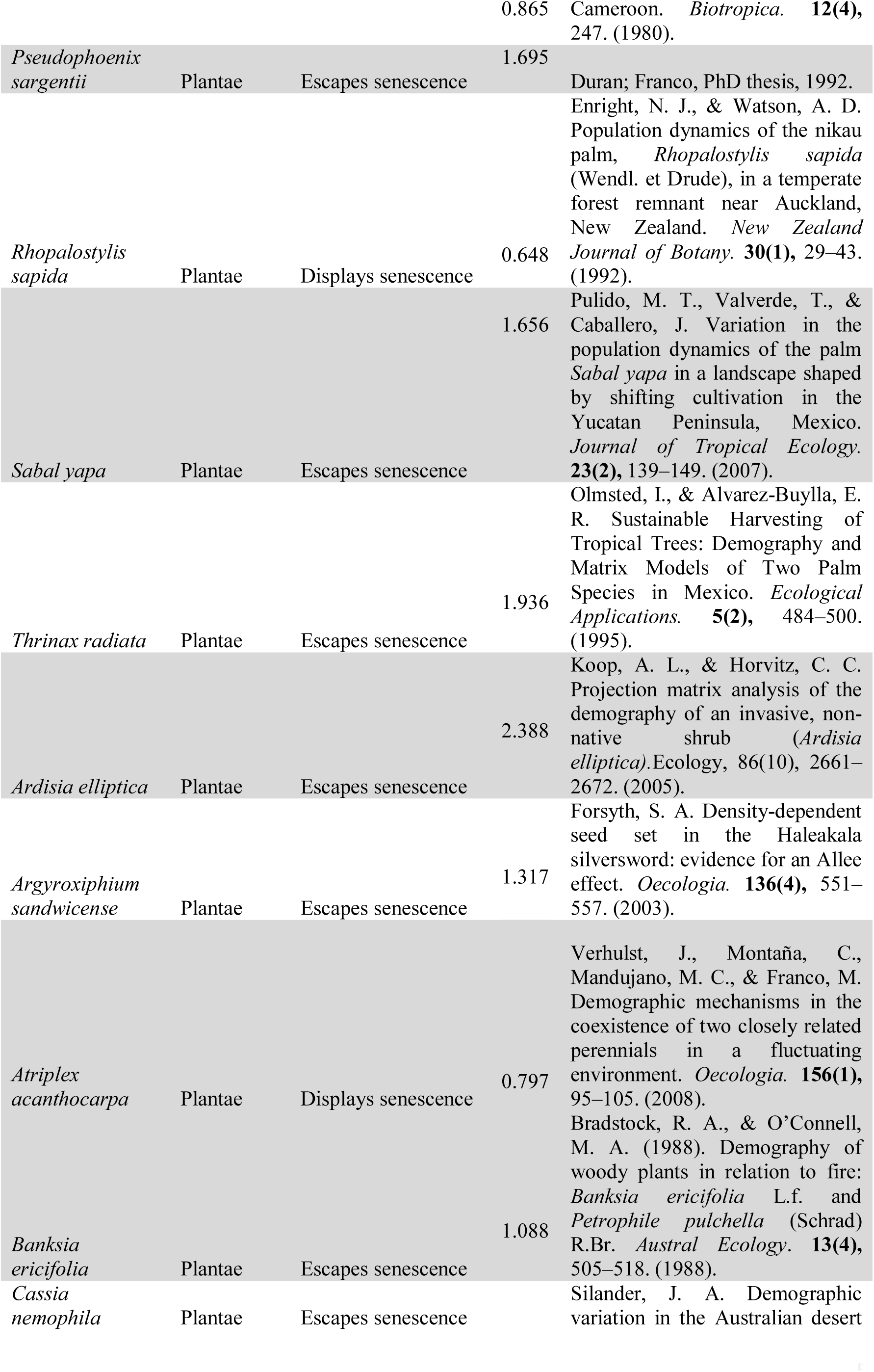
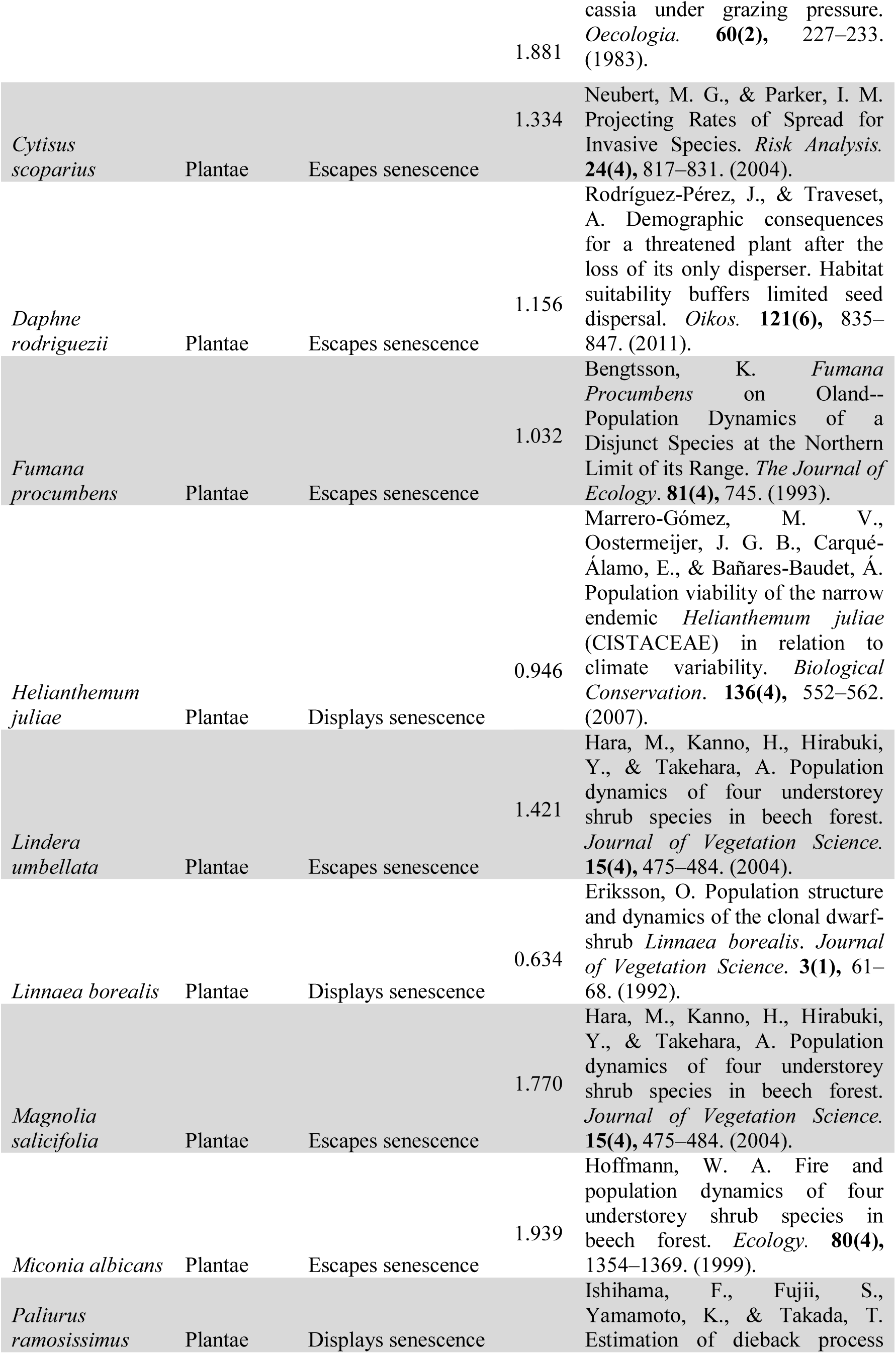
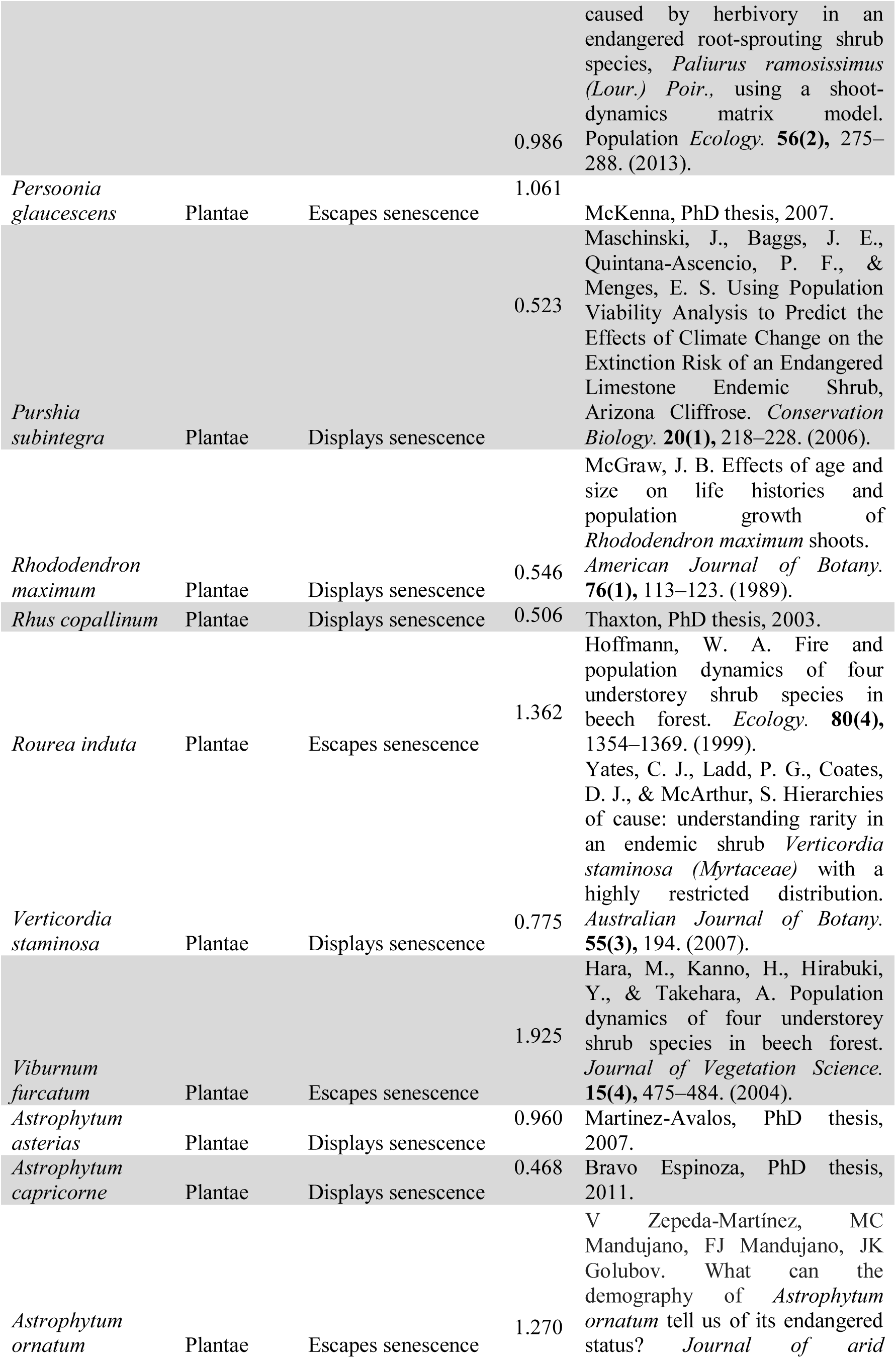
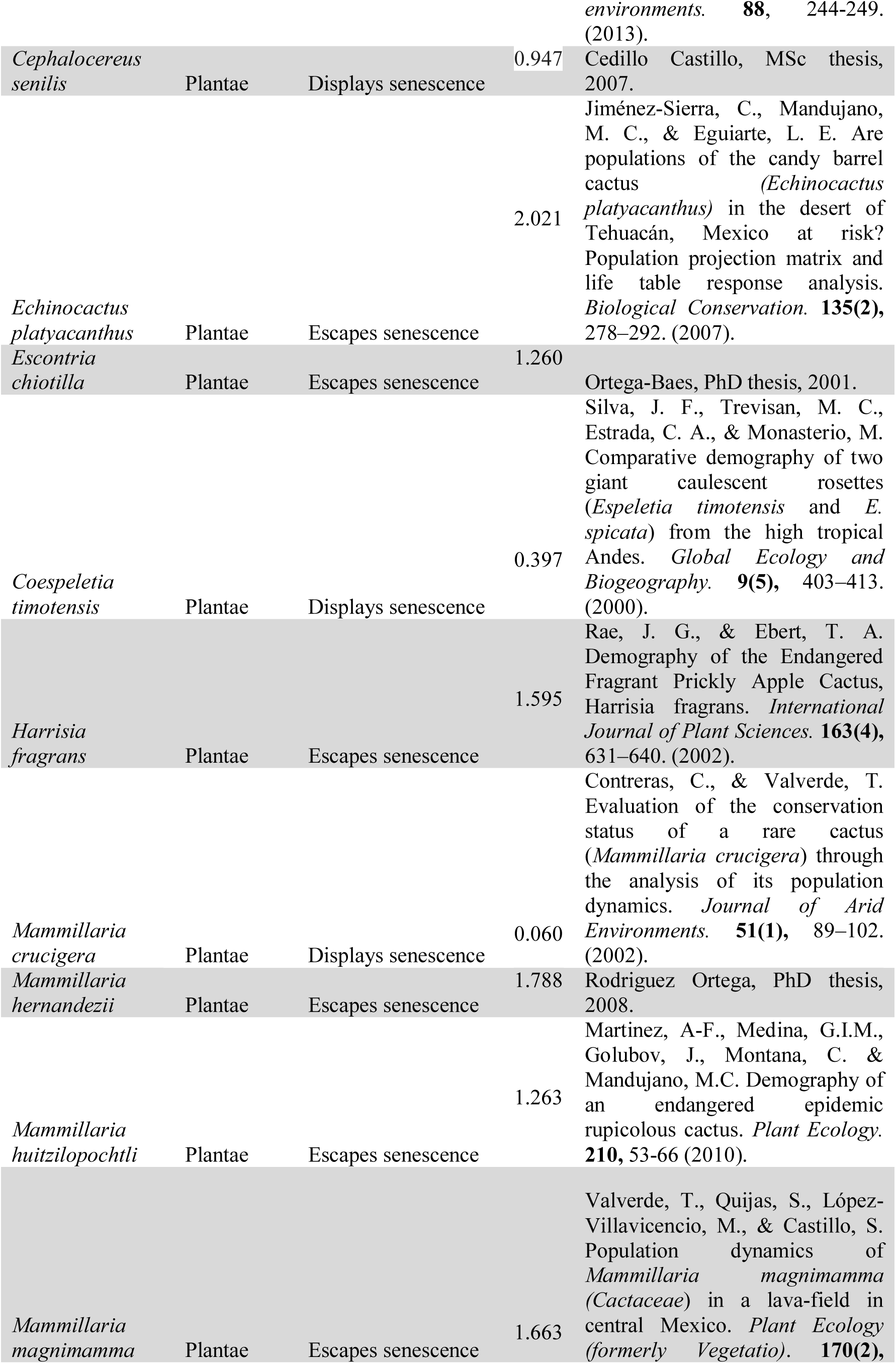
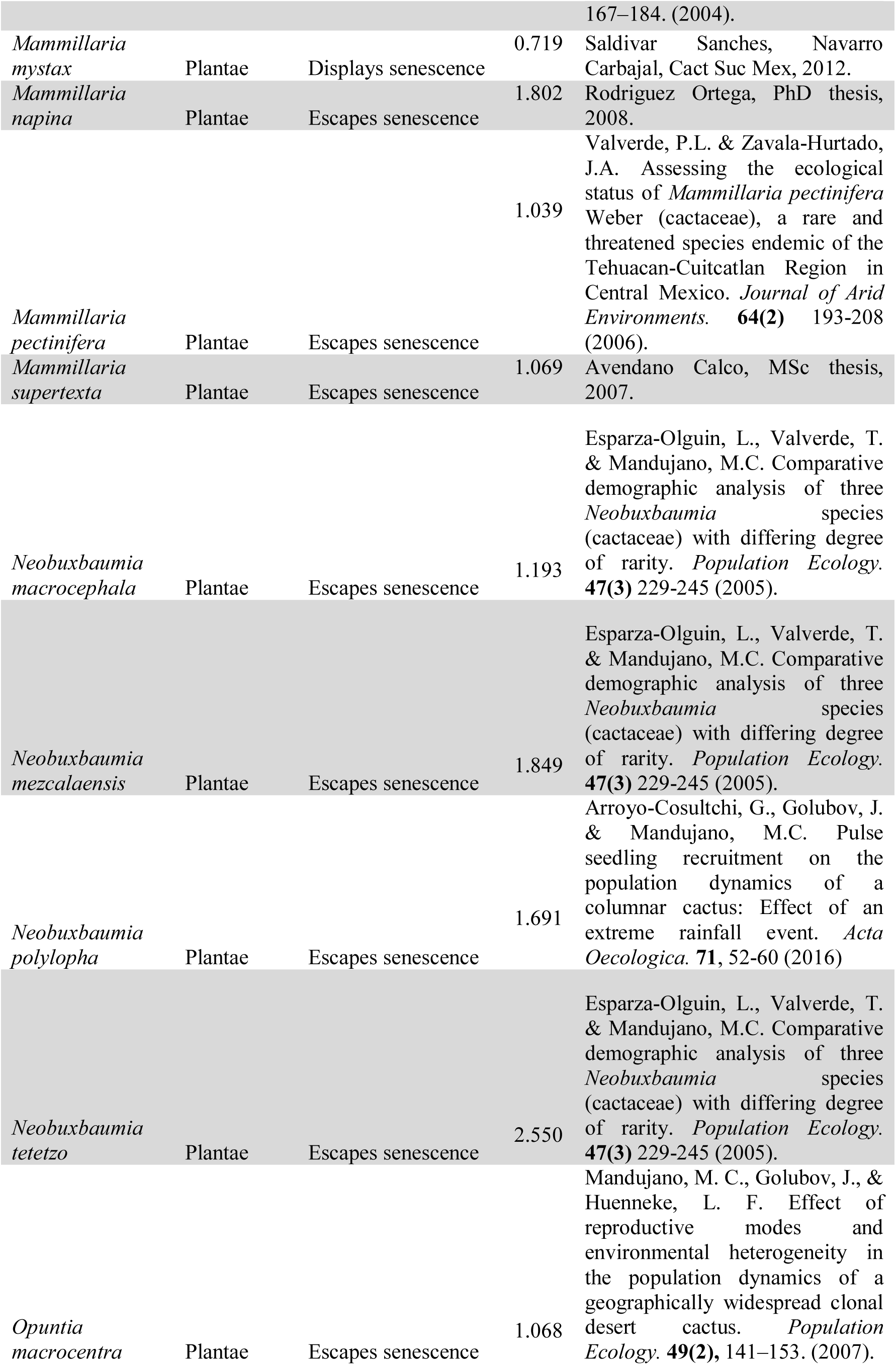
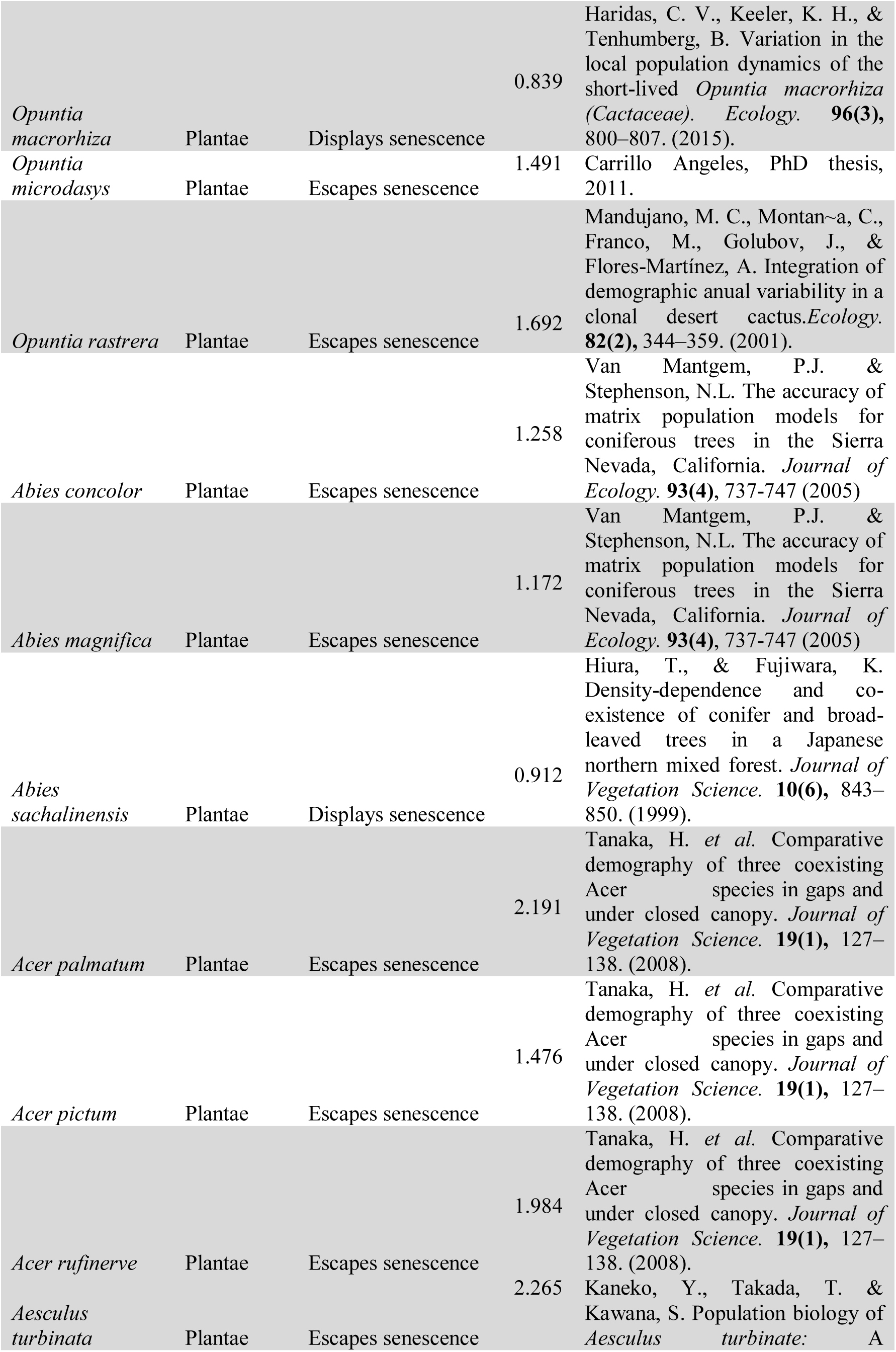
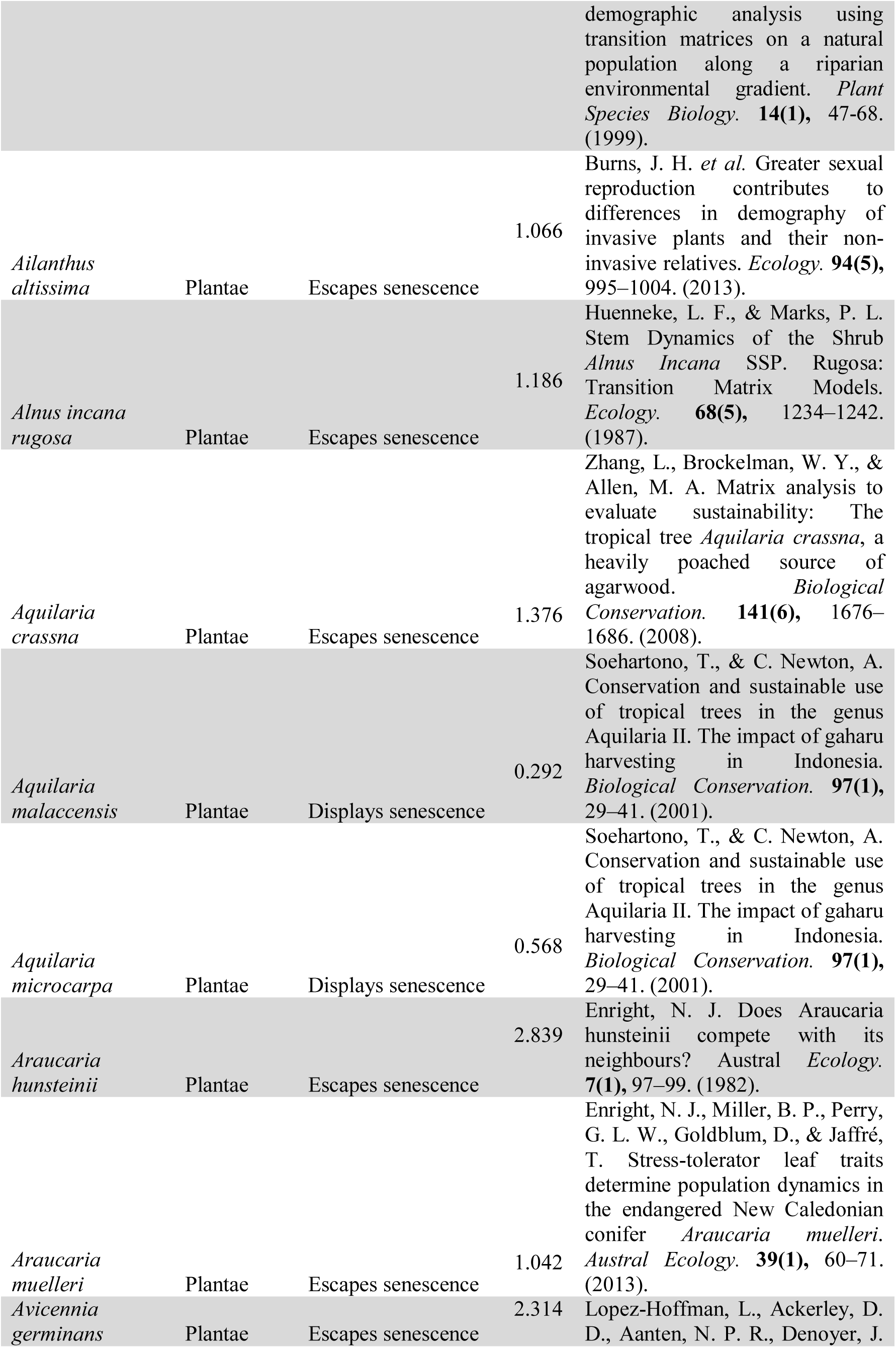
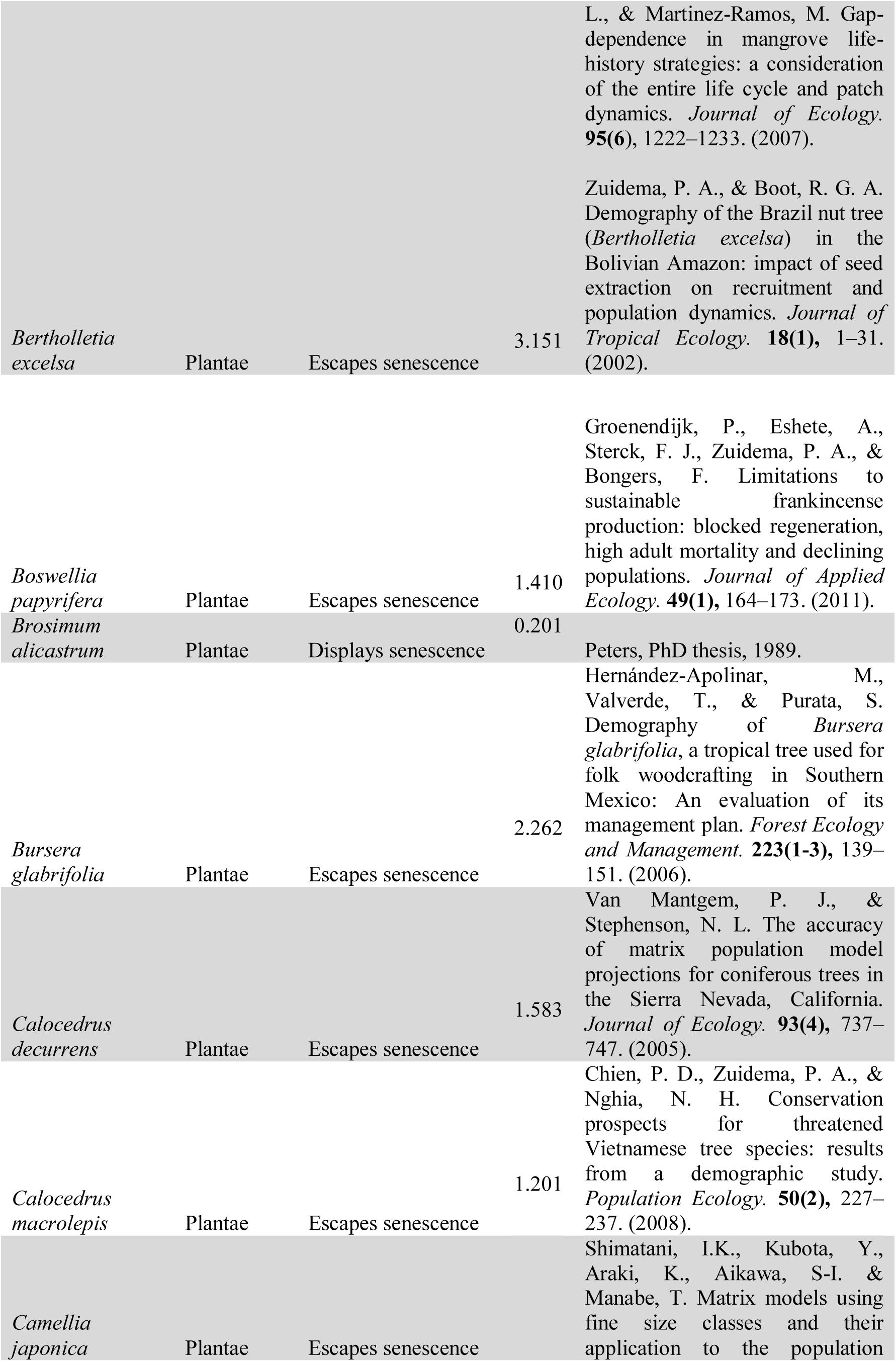
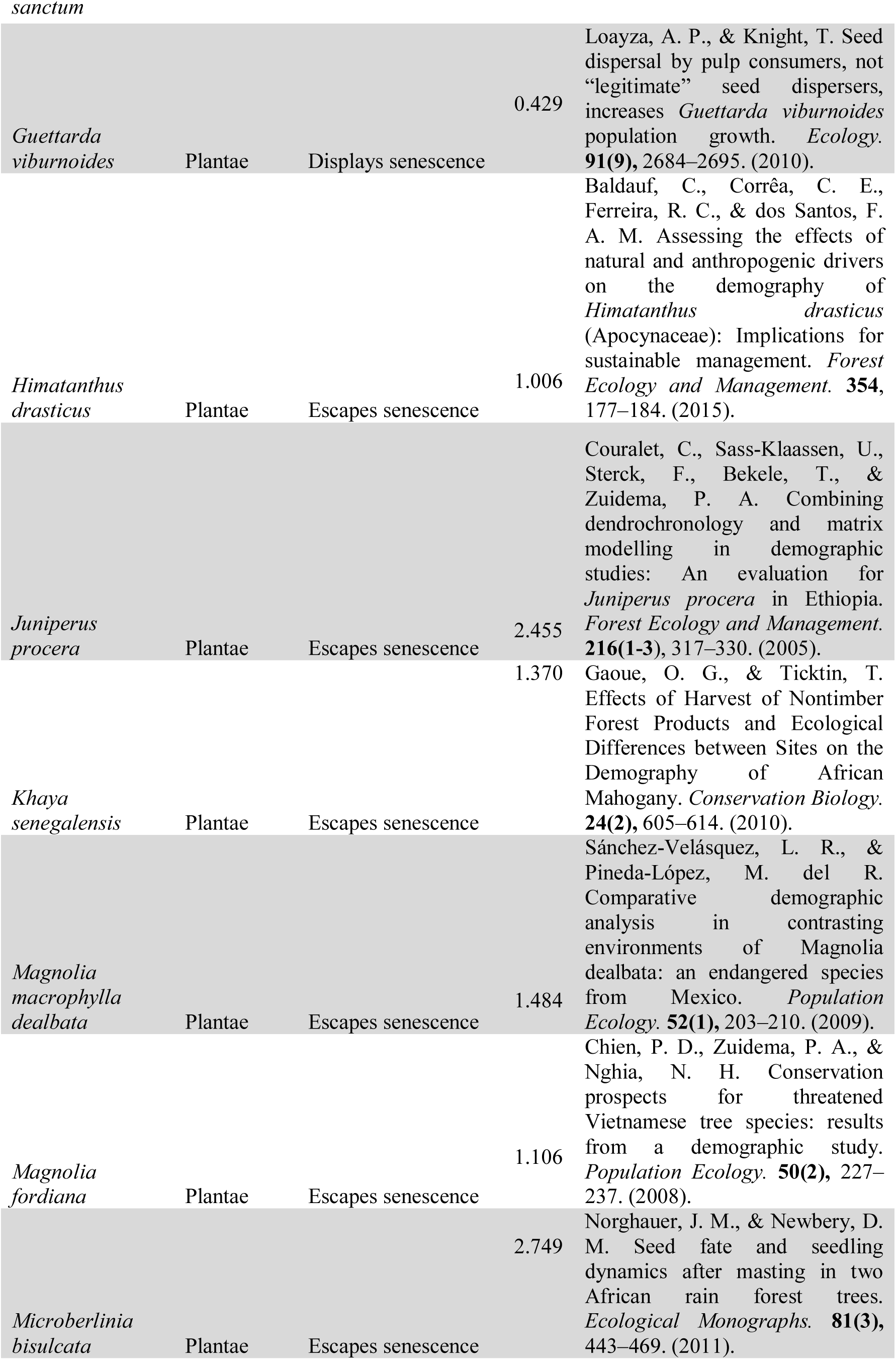
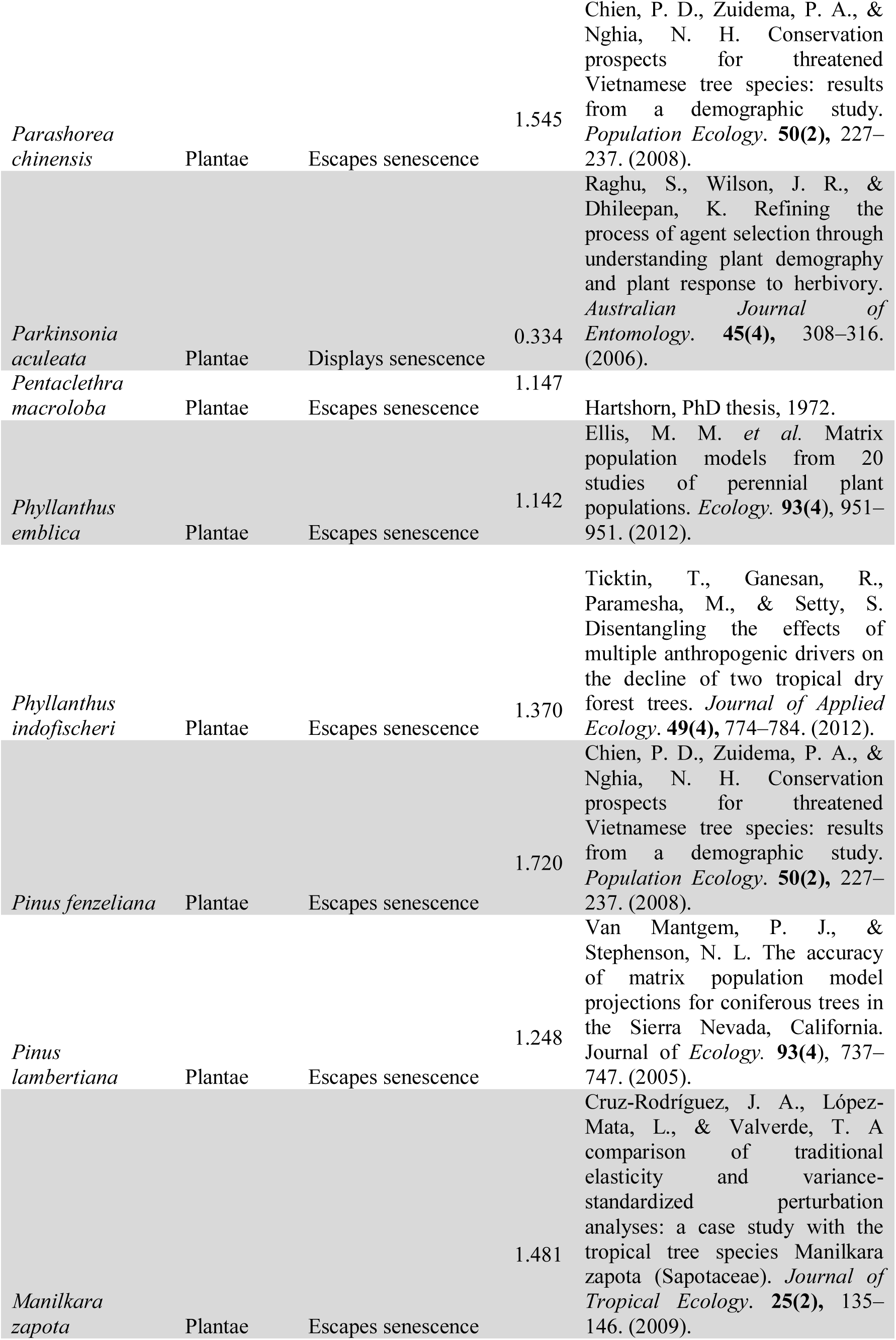
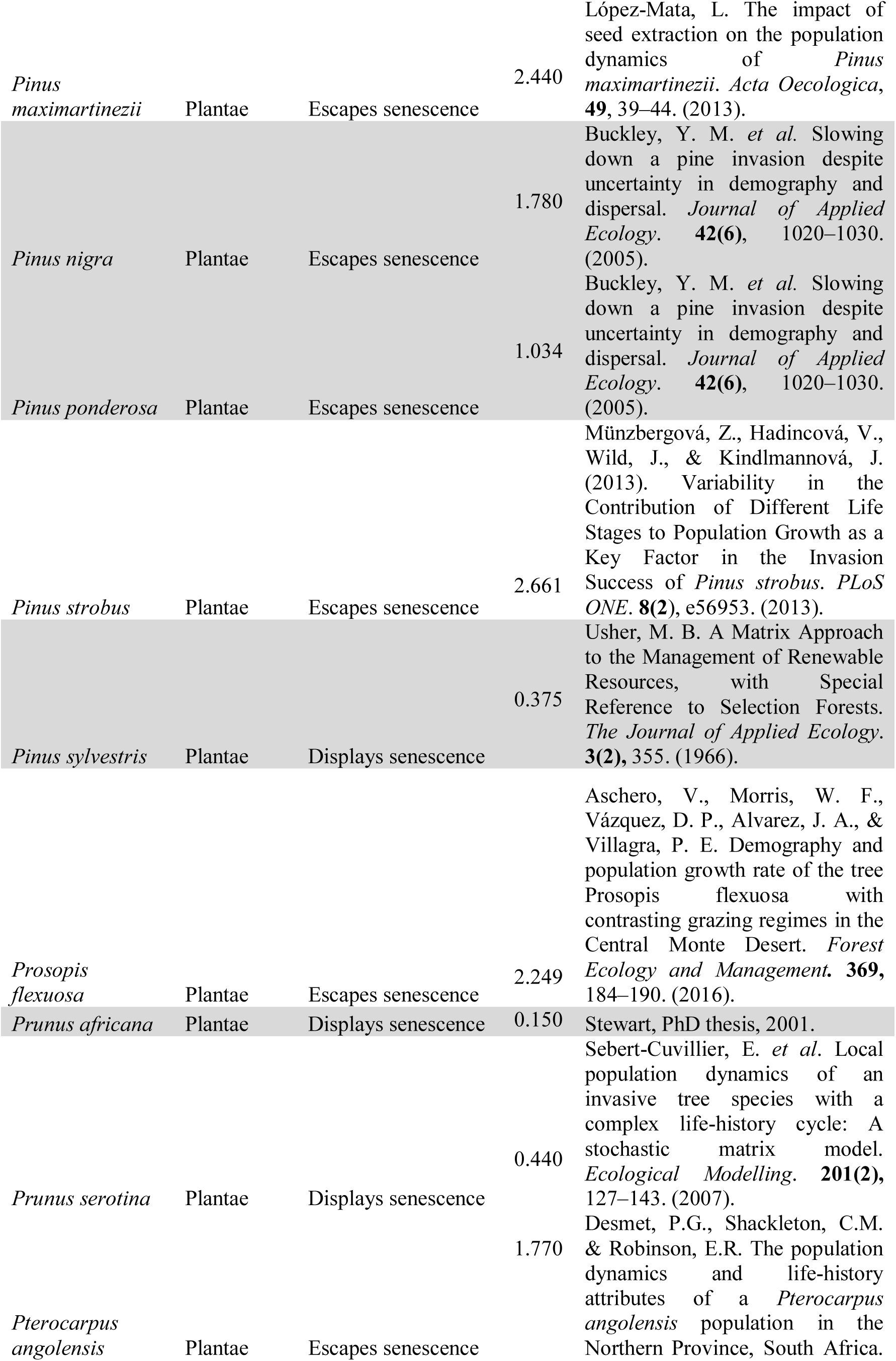
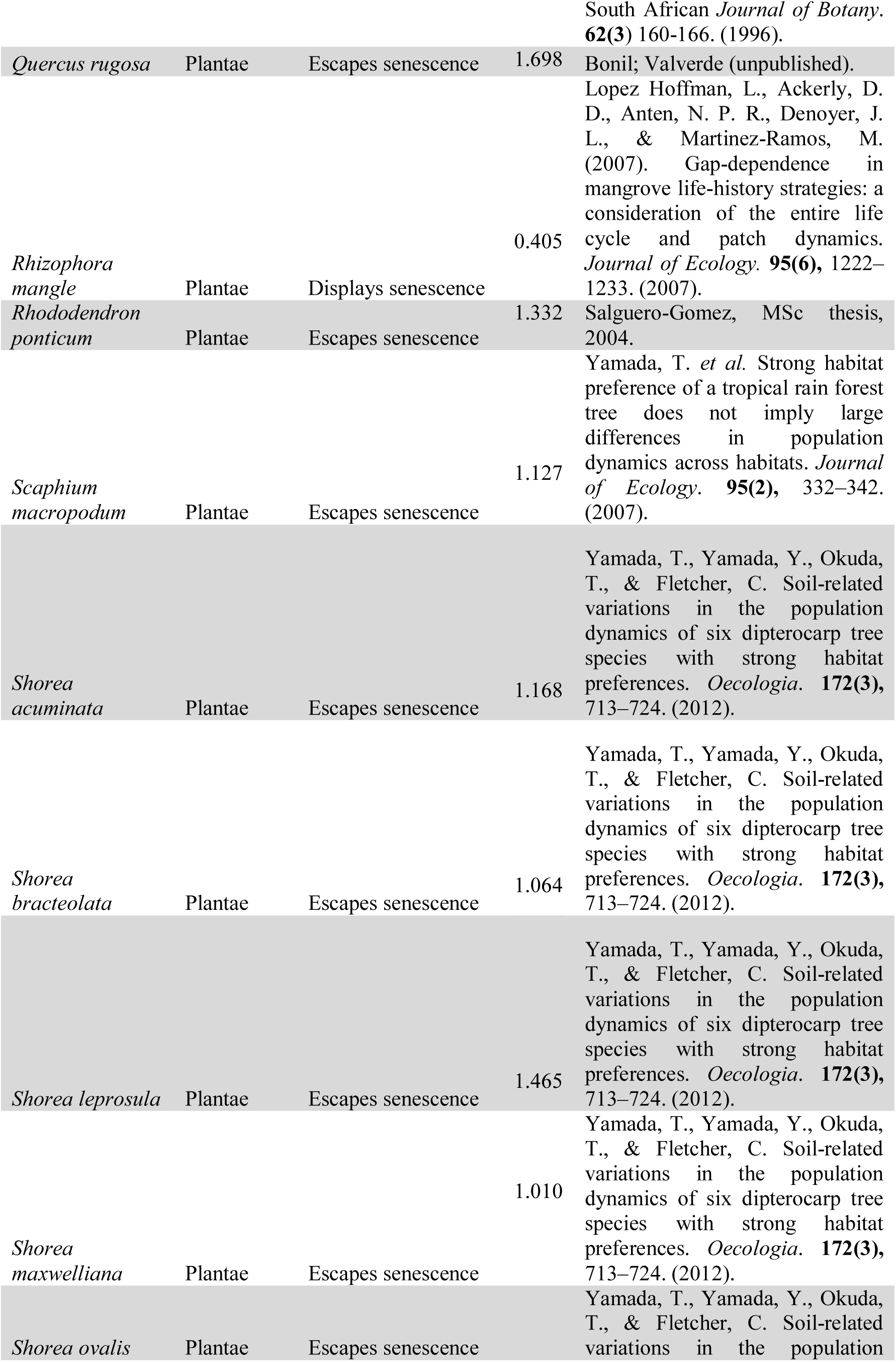
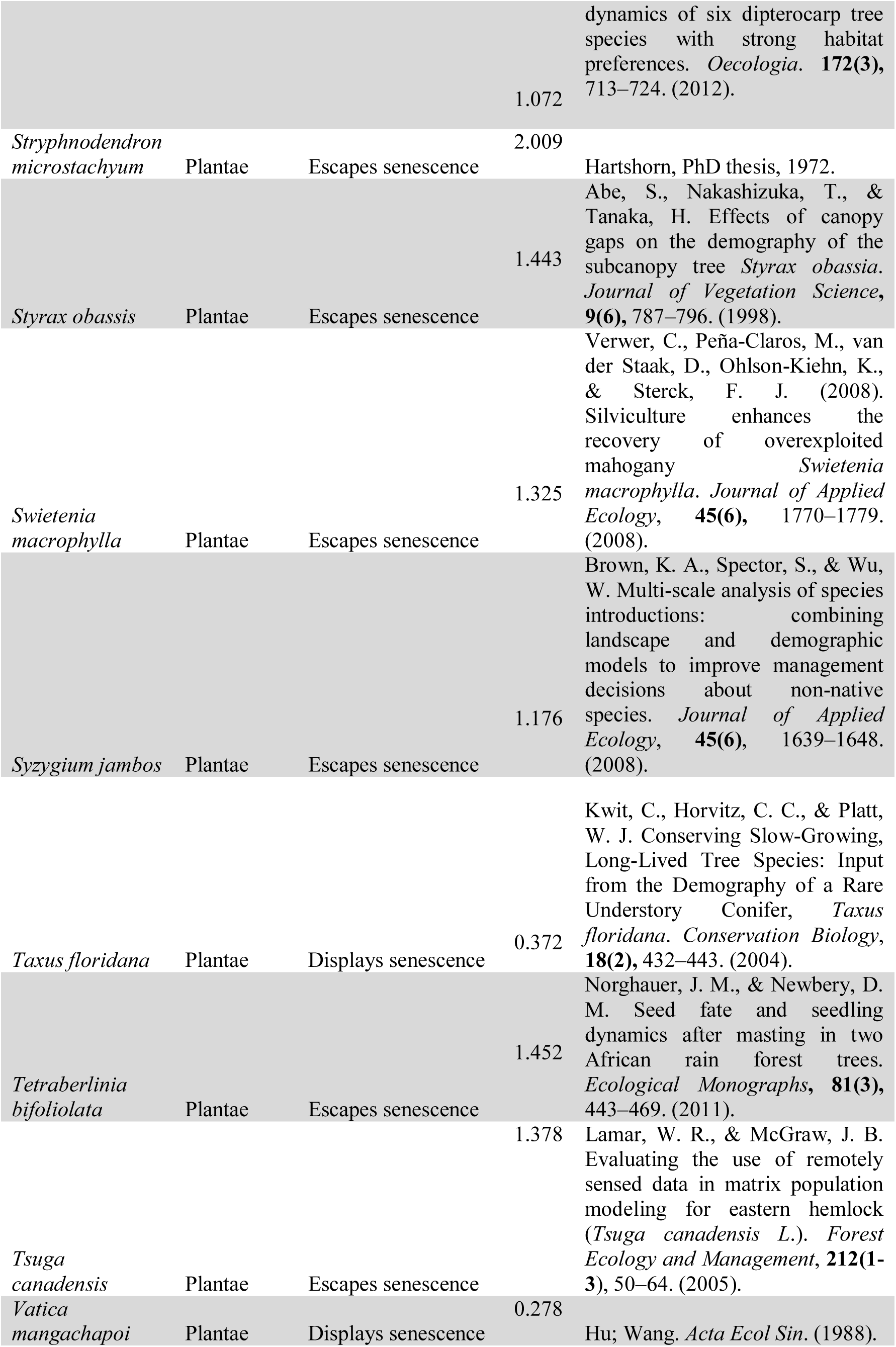
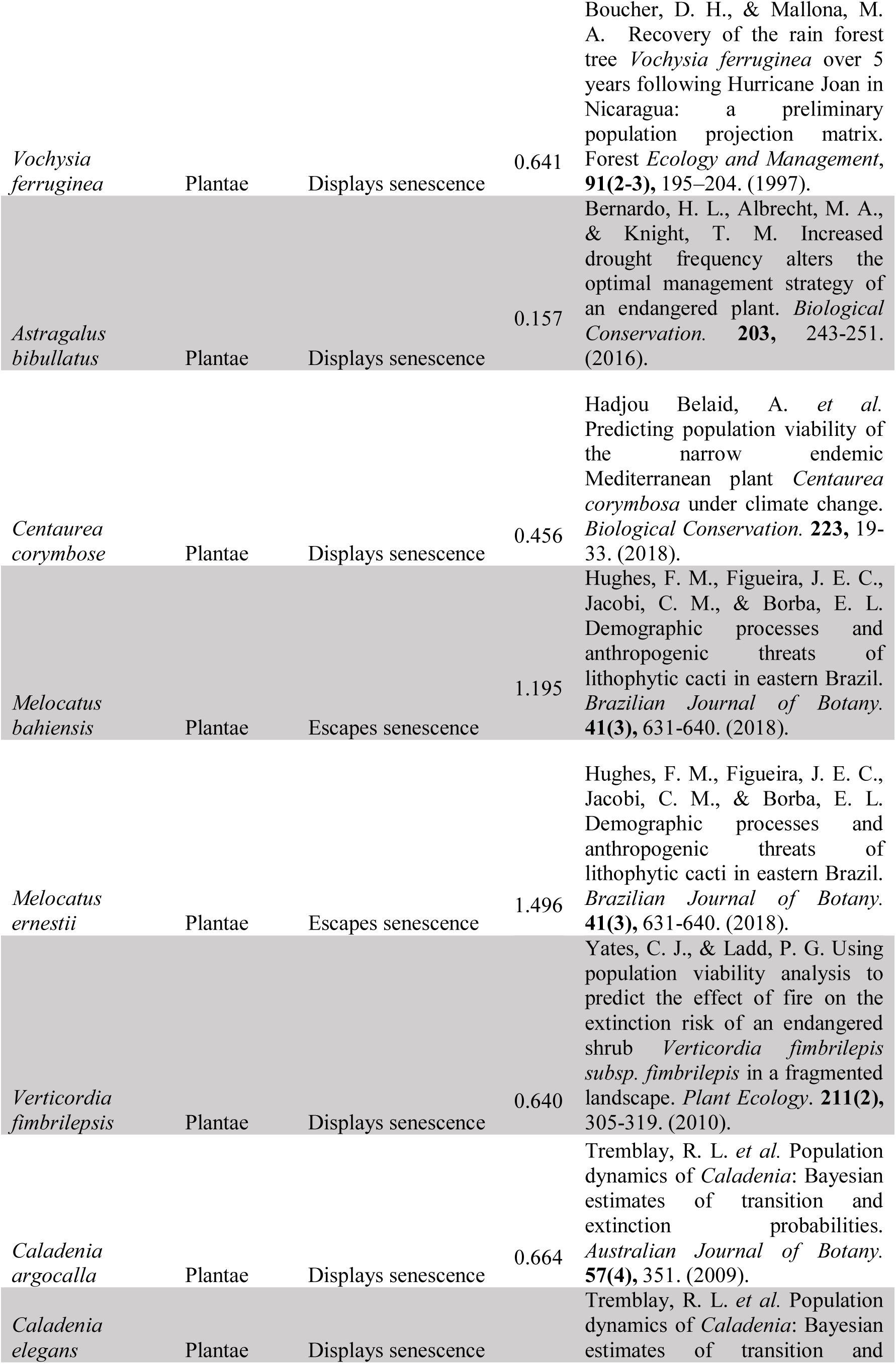
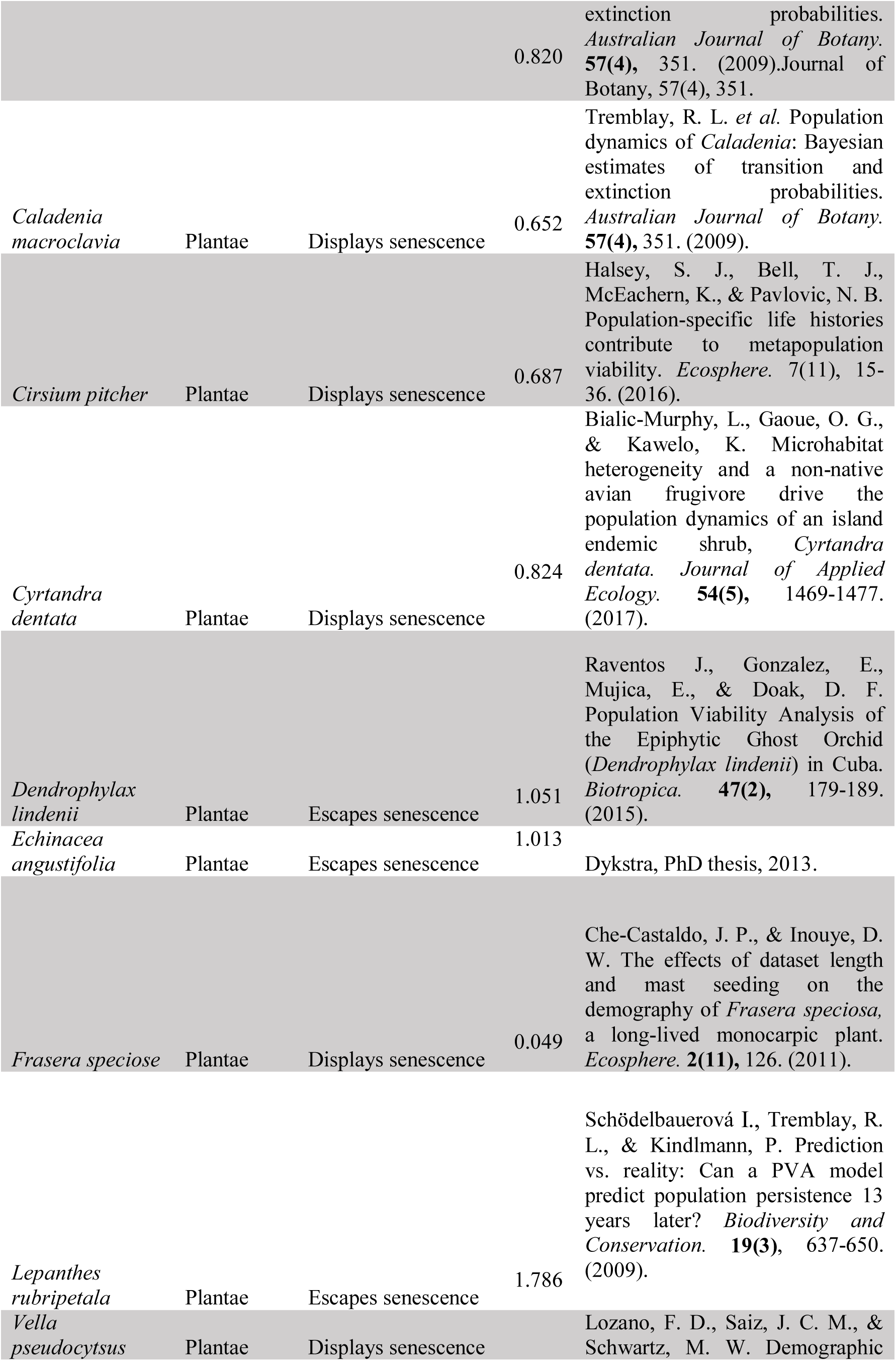
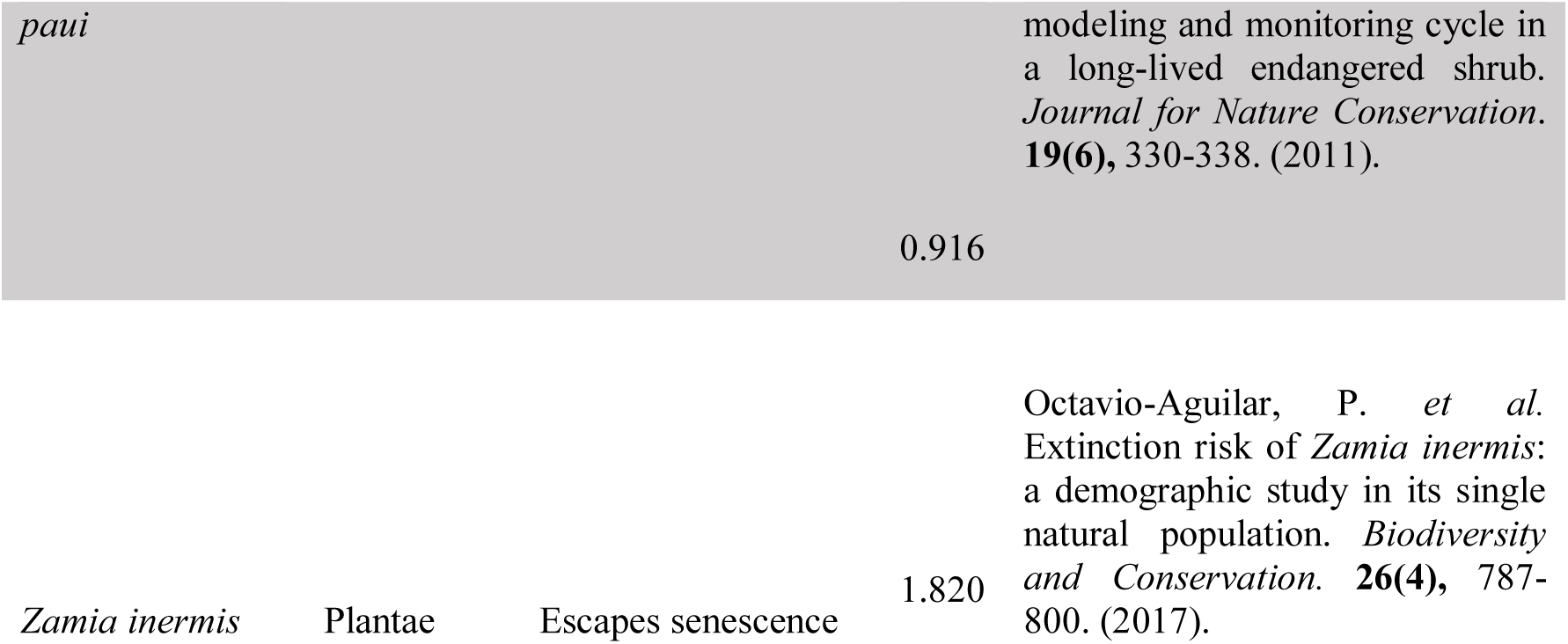
Study species. Included species in our analysis were those that fitted the selection criteria described in methods. Species names, their kingdom (Animalia or Plantae), whether they display or escape actuarial senescence according to Keyfitz’ entropy^14^ (*H),* and the source study from which the data was originally compiled into COMADRE^2^ and COMPADRE^3^ are listed.

**Supplementary Methods: ‘R’ computer code to extract age trajectories of fertility and mortality from population projection matrices, and to calculate Keyfitz’ entropy.**

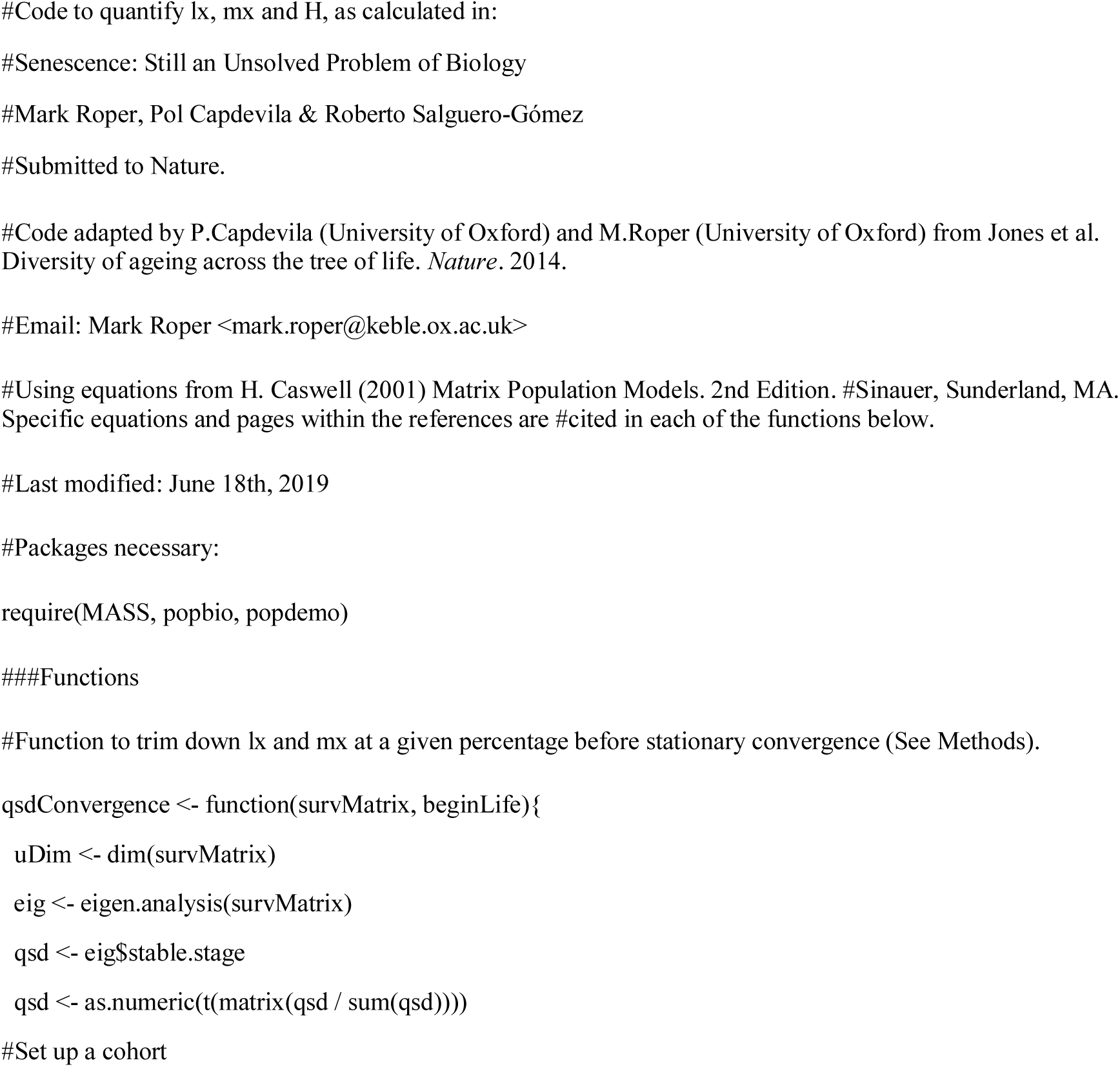
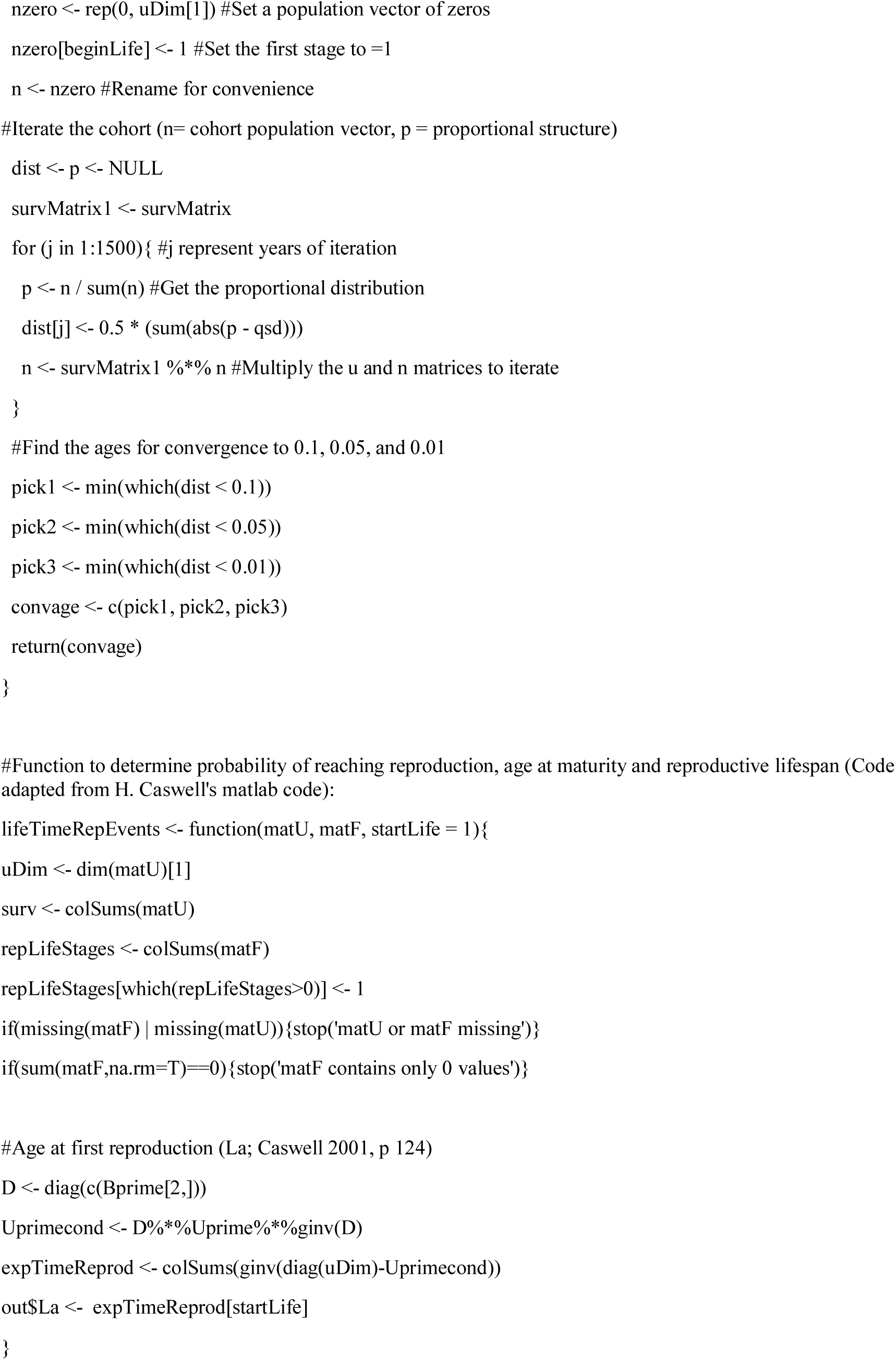
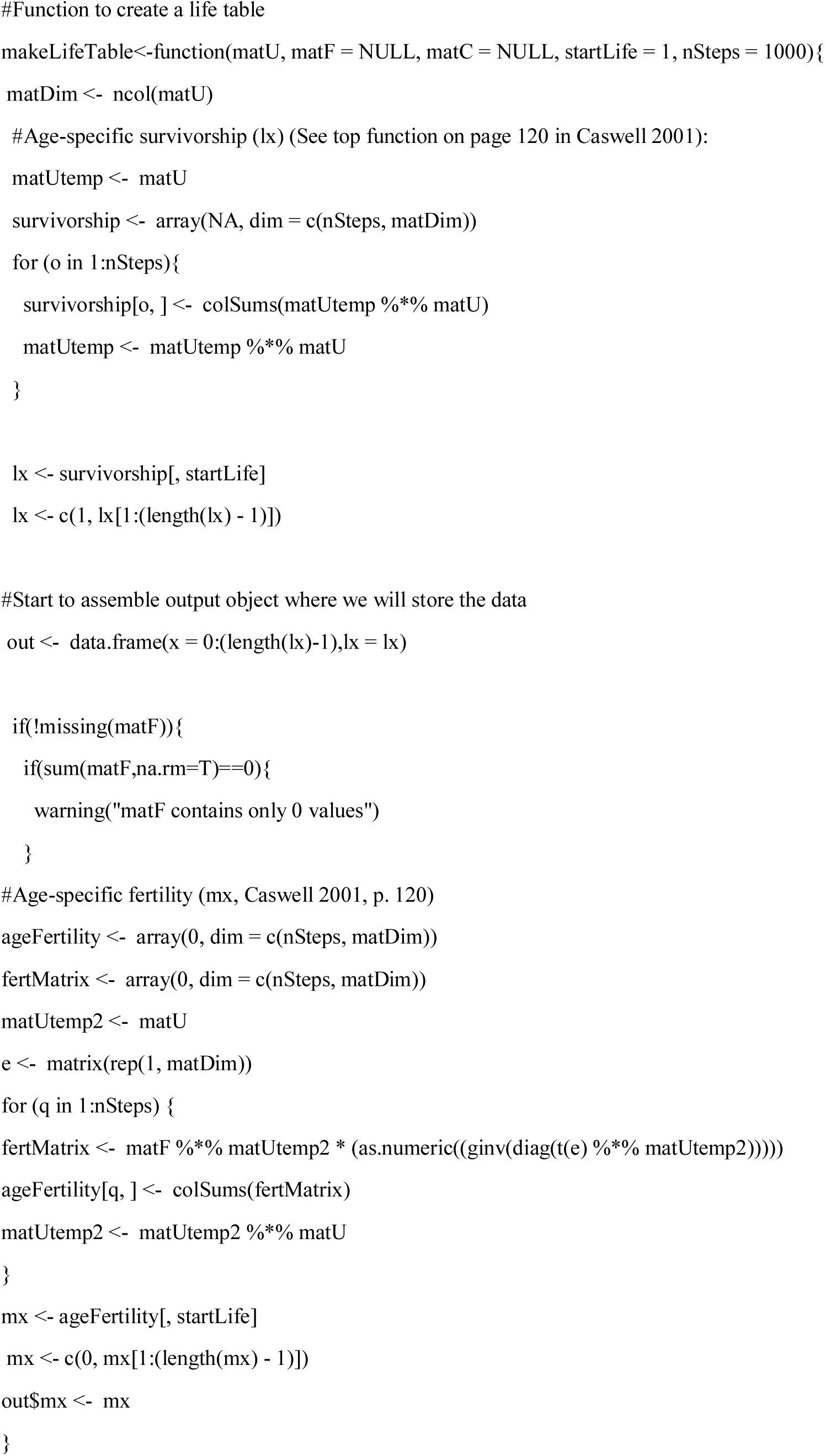
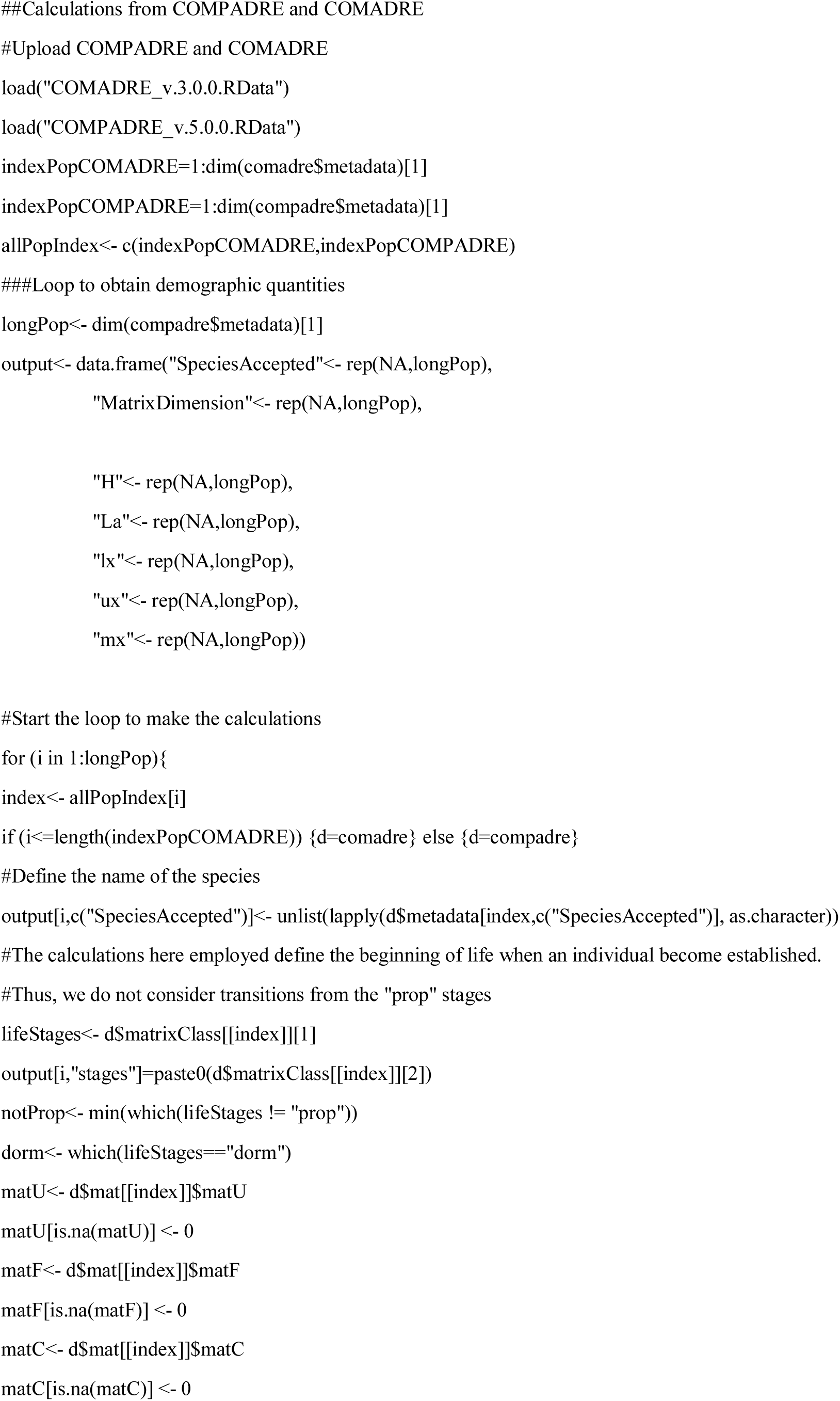
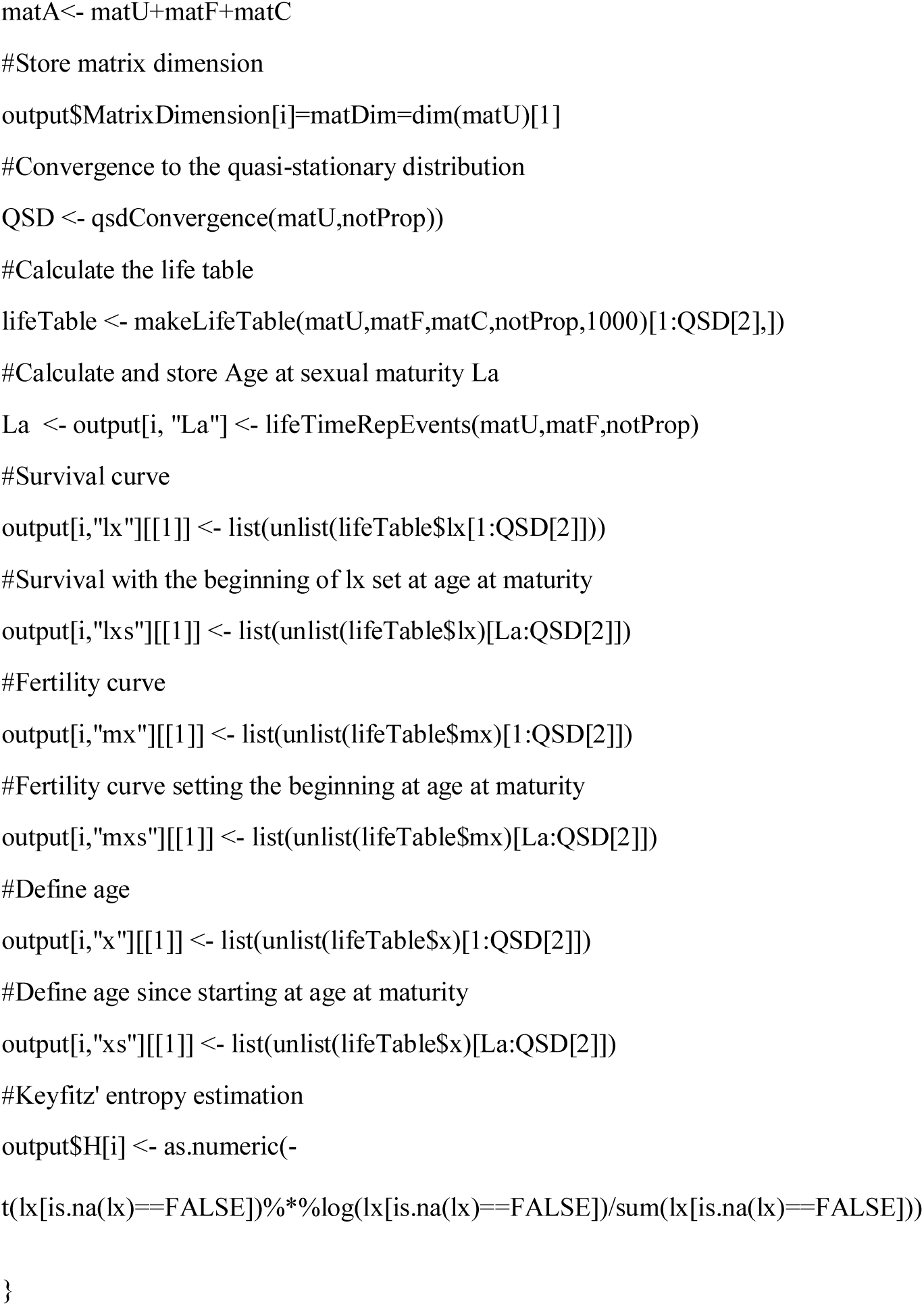

**Source data**

COMADRE_v.3.0.0 (as file)

COMPADRE_v.5.0.0 (as file)

